# Engineered 3D Hydrogel Matrices to Modulate Trophoblast Stem Cell-Derived Placental Organoid Phenotype

**DOI:** 10.1101/2024.05.13.594007

**Authors:** Emily M. Slaby, Nathaniel Hansen, Ritin Sharma, Patrick Pirrotte, Jessica D. Weaver

## Abstract

Placental organoid models are a promising platform to study human placental development and function. Organoid systems typically use naturally derived hydrogel extracellular matrices (ECM), resulting in batch-to-batch variability that limits experimental reproducibility. As an alternative, synthetic ECM-mimicking hydrogel matrices offer greater consistency and control over environmental cues. Here, we generated trophoblast stem cell-derived placental organoids using poly(ethylene glycol) (PEG) hydrogels with tunable degradability and placenta-derived ECM cues to evaluate trophoblast differentiation relative to Matrigel and two-dimensional (2D) culture controls. Our data demonstrate that PEG hydrogels support trophoblast viability and metabolic function comparable to gold standard Matrigel. Additionally, phenotypic characterization via proteomic analysis revealed that PEG and Matrigel matrices drive syncytiotrophoblast and extravillous trophoblast-dominant placental organoid phenotypes, respectively. Further, three-dimensional (3D) environments promoted greater integrin expression and ECM production than 2D culture. This study demonstrates that engineered 3D culture environments can be used to reliably generate placental organoids and guide trophoblast differentiation.

## INTRODUCTION

The human placenta is a vital organ during pregnancy that provides nutrients and protection to the developing fetus. The placenta is derived from the blastocyst’s outer layer, which differentiates into three subtypes of trophoblasts that form the complex placenta structure. Cytotrophoblasts (CT) are the proliferative progenitor to both the syncytiotrophoblasts (ST) and extravillous trophoblasts (EVT). STs form multinucleated, syncytialized cells that provide a barrier of protection and transport nutrients, oxygen, and waste for the fetus.^1^ EVTs invade into the maternal decidua during placental growth and aid in remodeling the vascular environment.^2^ The success of pregnancies relies on the healthy development, maintenance, and tight regulation of placenta functions; however, it is currently challenging to study the early stages of human placenta development due to suboptimal in vitro models.^3^ Furthermore, rodent models exhibit substantial physiological differences compared to the human placenta, limiting their utility.^4^ As such, in vitro models such as human placental organoids may bridge this gap to study early gestation trophoblast differentiation and subtype functions.

Recent models of human placental growth and differentiation use trophoblast stem cells (TSC), highly proliferative CTs derived from early gestation placentas that can differentiate into ST and EVT subtypes.^5–7^ TSCs are typically grown in two-dimensional (2D) systems as monolayers and in three-dimensional (3D) extracellular matrix (ECM)-rich Matrigel^TM^ with CT, ST, and EVT differentiation achieved using chemical signals;^8,9^ however, these models often fail to recapitulate the placental architecture and dynamic changes that occur as pregnancy progresses. Further, naturally derived matrices like Matrigel have high batch-to-batch variability in growth factors and ECM components. This variability may affect experimental reproducibility by influencing cell differentiation in unpredictable ways.^10^ As such, the development of 3D TSC culture models using highly defined synthetic materials could provide a solution to combat these issues.

ECM-mimicking synthetic hydrogel systems with tunable features and high biocompatibility offer promising alternatives to naturally derived matrices. The multi-arm poly(ethylene) glycol-maleimide (PEG-mal) system is one such popular system, as the maleimide functional groups rapidly react via Michael-type addition with free thiols, enabling functionalization with cysteine-terminated adhesive peptides and crosslinking with linear dithiol molecules. This system has been used to culture primary islet organoids,^11–13^ intestinal organoids,^14^ and mesenchymal stem cells,^15,16^ among others.^17–21^ We previously employed a PEG-mal hydrogel system with tunable degradability and ECM-derived adhesive ligand sequences to culture trophoblast-like choriocarcinoma cell lines;^22^ however, choriocarcinoma cell lines are a poor model to recapitulate the placenta architecture and trophoblast subtypes.

Here, we use primary TSCs with a proliferative CT phenotype that can be differentiated into ST and EVT subtypes by established protocols^5–7^ to study the effects of synthetic and natural hydrogel culture on placental organoid growth and differentiation. We hypothesized that a 3D synthetic hydrogel crosslinked with protease-sensitive linkers and modified with ECM-derived adhesive ligands present in the placenta would support trophoblast viability and differentiation comparably to gold-standard Matrigel. We demonstrate that synthetic PEG hydrogels support the viability and differentiation of placental organoids into a predominantly ST-like phenotype, while Matrigel culture environments result in a predominantly EVT-like phenotype. This synthetic hydrogel culture system may enhance the modeling of early placenta development, with less lot-to-lot variability than natural ECM matrices, with the potential to facilitate drug discovery and advance therapeutic interventions in obstetric care.

## METHODS

### Materials

Crosslinker and adhesion ligand sequences were purchased from GenScript as outlined in Table 1, except for crosslinkers DTT (ThermoFisher catalog #R0861) and PEG-dithiol (DT, Mn = 1000, Sigma catalog #717142; Mn = 3000, Sigma catalog #704539). Cell culture reagents were purchased from Fisher Scientific unless otherwise noted.

**Table 1.** Crosslinker and adhesion ligand peptide amino acid sequences.

| Name | Amino Acid Sequence | Description of Function within Matrix |
| --- | --- | --- |
| RGD | GRGDSPC | Adhesive ligand, fibronectin |
| RDG | GRDGSPC | Adhesive ligand, scrambled fibronectin |
| GFOGER | GGYGGGPP(GPP)5GFOGER(GPP)5GPC | Adhesive ligand, collagen |
| YIGSR | CGGEGYGEYIGSR | Adhesive ligand, laminin |
| IKVAV | CGQAASIKVAVSADR | Adhesive ligand, laminin |
| VPM | GCRDVPMSMRGGDRCG | Crosslinker, degradable by MMP-1, MMP-2, MMP-3, MMP-7, MMP-9, MT1-MMP/MMP-14<sup>23-25</sup> |
| GDQ | GCRDGDQGIAGFDRCG | Crosslinker, degradable by MMP-1<sup>22,24,26,27</sup> |
| GPQ-W | GCRDQGWIGQPGDRCG | Crosslinker, degradable by MMP-1, MMP-2, MMP-3, MMP-8, MMP9, MMP-12<sup>24,28,29</sup> |

### Trophoblast Stem Cell (TSC) Culture

Human TSC CT 1049, derived from 6-week 1-day gestation age male fetus placental tissues,^5^ were kindly gifted by Dr. Mana Parast from the University of California, San Diego. These cells were cultured with modifications to previously published literature^6^ on 5 µg/mL collagen-coated (Sigma catalog #C0543-1VL, incubated >90 minutes in PBS at 37 °C) 6-well plates at a density of 0.2-0.5 x 10^6^ cells per well in advanced DMEM/F12 (Gibco^TM^ catalog #12634010), 1X B27 (Gibco catalog #17504044), 1X N2 (Gibco catalog #17502048), 1X GlutaMAX (Gibco catalog #35050061), 150 µM 1-thioglycerol (Sigma catalog #M6145), 1% KnockOut Serum Replacement (Gibco catalog #10-828-028), 0.05% bovine serum albumin (BSA, Gemini Bio Products catalog #50-753-3079), 2 µM CHIR99021 (Sigma catalog #SML1046), 500 nM A83-01 (Tocris Bioscience^TM^ catalog #29-391-0), 1µM SB43154 (Sigma catalog #616464), 0.8 mM valproic acid sodium salt (Sigma catalog #676380), 5 µM Y27632 (Selleck catalog #S1049), 100 ng/mL FGF2 (R&D Systems catalog #3718FB01M), 50 ng/mL EGF (R&D Systems catalog #236-EG-01M), 50 ng/mL HGF (Stem Cell Technologies catalog #78019), and 20 ng/mL Noggin (R&D Systems catalog #6057-NG) with media replenished every 2-3 days or passaged with TrypLE (Gibco catalog #1260421) at 60-80% confluence. Cells were maintained in a humidified incubator at 37 °C in 5% CO_2_.

### ST and EVT Differentiation

ST differentiation was achieved with modifications to previously published literature^6^ by culturing passaged CT cells on 5 µg/mL collagen-coated (Sigma catalog #C0543-1VL, incubated >90 minutes in PBS at 37 °C) 6- or 24-well plates (2D conditions only) in advanced DMEM/F12, 1X B27, 1X N2, 4% KnockOut Serum Replacement, 0.05% BSA, 2.5 mM Y27632, and 2 mM forskolin (Sigma catalog #344270) with media replenished every 2 days.

EVT differentiation was achieved with modifications to previously published literature^6^ by culturing passaged CT cells on 20 µg/mL fibronectin-coated (Gibco catalog #PHE0023, incubated >90 minutes in DPBS at 37 °C) 6- or 24-well plates (2D conditions only) in advanced DMEM/F12, 0.1 mM 2-mercaptoethanol (Gibco catalog #21985023), 0.3% BSA, 1% ITS-X supplement (Gibco catalog #51500-056), 7.5 µM A83-01, 4% KnockOut Serum Replacement, 100 ng/mL NRG1 (Cell Signaling catalog #26941), and 2% Matrigel (Corning catalog #354234) for 4 days (changed every 2 days), then changing the media to 0.5% Matrigel without NRG1 for another 2 days. Cells were maintained in a humidified incubator at 37 °C in 5% CO_2_.

### Flow Cytometry

CTs were plated on coated 6-well plates for 6 days with CT, ST, or EVT media changes on days 2 and 4 following the methods above. On day 6, cells were washed with DPBS and lifted with TrypLE, then washed in DPBS + 1% bovine serum albumin (BSA). Human TruStain FcX^TM^ (FC Receptor Blocking Solution, BioLegend catalog #422302) was incubated with the cells for 5 minutes at room temperature at a 1:250 dilution in DPBS with 1% BSA. Cells were then stained with Brilliant Violet (BV) 421^TM^ anti-human CD138 (Syndecan-1) antibody (BioLegend catalog #356516, clone MI15), FITC anti-human HLA-A, B, C antibody (BioLegend catalog #311404, clone W6/32), and APC anti-human EGFR antibody (BioLegend catalog #352906, clone AY13) at 1:100 dilutions in DPBS + 1% BSA for 15 minutes at room temperature in the dark. Cells were washed with DPS + 1% BSA. Cells were stained with stained for viability with Zombie Aqua^TM^ Fixable Viability Kit (BioLegend catalog #423102) at 1:1000 in DPBS for 30 minutes at room temperature following the manufacturer’s instructions, then washed with DPBS with 1% BSA. Cells were fixed with 4% paraformaldehyde for 20 minutes at room temperature, then washed with DPBS + 1% BSA and stored in DPBS + 1% BSA at 4 °C until analysis on a Life Technologies Attune NxT with Autosampler. Samples were analyzed in FlowJo^TM^ 10, gating out debris and gating on single and live events.

### Hydrogel Preparation

CT cells were passaged at 60-80% confluence with TrypLE in standard 2D culture conditions and encapsulated in 3D hydrogel matrices for 6 days maintained in CT media or switched to ST or EVT differentiation medium. PEG-mal (20 kDa, 4-arm, 5% w/v, Laysan Bio catalog #4arm-PEG-MAL-20K) macromer with 1 mM adhesion ligand (RGD, RDG, GFOGER, YIGSR, or IKVAV, see Table 1) and protease-degradable (VPM, GDQ, GPQ-W, see Table 1) and/or nondegradable (DTT, PEG-DT: MW = 1000 Da or 3400 Da) was used to encapsulate cells as previously described.^22,30^ Briefly, PEG-mal, adhesion ligands, and crosslinker reagents were resuspended in DPBS, and the pH was adjusted to 7, if necessary, with sodium hydroxide. Matrigel (Corning catalog #356237) was used at 5 mg/mL (0.5% w/v), diluted with cell-specific media, while placenta-derived human collagen IV (Corning catalog #354245) was used as provided. Low-gelling temperature agarose (Sigma catalog #A9045) was resuspended in DPBS (with magnesium and calcium chloride) to acquire a 2% w/v solution and heated to 100 °C for 30 minutes with stirring for homogenization. Sodium alginate (120 mM, 4% w/v; Novamatrix catalog #4200101) and 20 mM glucono-delta-lactone (GDL) were resuspended in DPBS and incubated at 4 °C for >12 hours for homogenization. Calcium carbonate (60 mM) was added to the sodium alginate solution before use and mixed at a 1:1 ratio with the GDL solution to acquire a final 2% sodium alginate hydrogel. CTs maintained in CT media were encapsulated with 100,000 cells per 15 µL hydrogel, while cells switched to ST or EVT differentiation medium were encapsulated with 200,000 cells per 15 µL hydrogel. 2D conditions were plated with the same number of cells per subtype condition into a 24-well culture-treated plate. Cell-specific media (1 mL) was added to the hydrogels or 2D wells in a 24-well plate and replaced on days 2, 4, and 6 based on the culture procedure above.

### Immunocytochemistry

After six days of culture, hydrogels were fixed with 4% paraformaldehyde (Thermo Scientific catalog #AAJ19943K2) at room temperature for 20 minutes, then stored in DPBS at 4 °C until staining. Hydrogels were incubated in DPBS with 1% BSA and 0.3% Triton-X100 for 60 minutes, then blocked with PowerBlock^TM^ Universal Blocking Reagent (BioGenex Laboratories catalog #HK085-5K) for 90 minutes at room temperature, then goat serum (BioGenex Laboratories catalog #HK1129K) for 30 minutes at room temperature. Hydrogels were stained with primary antibodies (mouse anti-KRT7, 1:300, NSJ Bioreagents catalog #V2656; rabbit anti-Ki67, 1:400, Cell Signaling catalog #9129; rat anti-E-cadherin, 1:250, Novus catalog #NB120-11512; mouse anti-syndecan-1, 1:500, biorbyt catalog #orb153547; rabbit anti-HLA-G, 1:300, Cell Signaling catalog #79769, and/or mouse anti-ZO-1-AF594, 1:500, ThermoFisher catalog #339194) or isotype controls (mouse IgG1 kappa, Novus catalog #NBP1-43319; rabbit IgG, Novus catalog #NBP2-24891; rat IgG1 lambda, Novus catalog #NBP2-31380; mouse IgG1, Novus catalog #NBP1-97005) diluted in DPBS with BSA and Triton-X100 for 2 hours at room temperature or overnight at 4°C. Hydrogels were washed three times in DPBS with BSA and Triton-X100 for 10 minutes. Secondary antibodies (AF488 goat anti-mouse IgG, Abcam catalog #ab150113; AF594 goat anti-rabbit, Abcam catalog #ab150080; AF680 goat anti-rat IgG, Abcam catalog #ab175778) were diluted 1:500 in DPBS with BSA and Triton-X100 and incubated for 75 minutes at room temperature in the dark. Hydrogels were washed thrice in DPBS with BSA and Triton-X100 for 10 minutes. Hydrogels were transferred to slides with ProLong^TM^ Gold Antifade Mountant with DAPI (Invitrogen catalog #P36931) and a coverslip to cure for 24 hours in the dark at room temperature before being stored at 4 °C until imaging on a Leica SP8 White Light Laser Confocal Microscope. Images were analyzed with Fiji using the same threshold, size, and circularity constraints.^31^

### Western Blotting

CTs were cultured in Matrigel for 6 days with CT, ST, or EVT differentiation media. On day 6, hydrogels were washed with DPBS and lysed in 1X RIPA Lysis Buffer (Millipore catalog #20-188) diluted in DI water with 1:50 Halt^TM^ Protease & Phosphatase Inhibitor Cocktail (catalog #1861284) for 10 minutes on ice and subsequently centrifuged at 13,000 x gravity for 10 minutes at 4 °C for protein lysate collection. Samples were stored at −80 °C until the electrophoresis run. Pierce^TM^ BCA (bicinchoninic acid) Protein Assay Kit (catalog #23225) was used to quantify protein concentration. Protein (10 μg) from each sample was combined with 1X Bolt LDS Sample Buffer (1:4, Invitrogen catalog #B0007) and 1X Bolt Reducing Agent (1:10, Invitrogen catalog #B0009) according to the manufacturer’s instructions. Samples were incubated at 100 °C for 10 minutes to denature proteins. Electrophoresis was performed in NuPAGE MES SDS Running Buffer (Invitrogen catalog #NP0002) on Bolt BisTris 4-12% 1.00 mm Mini gels (Invitrogen catalog #NW04120BOX) for 32 min at 200 V. The proteins were transferred using iBlot Transfer stacks (Invitrogen catalog #IB23001) to a nitrocellulose membrane with a 0.2 μm pore size using the iBlot 2 P0 preset method for 1 minute at 20 V, 4 minutes at 23 V, and 2 minutes at 25 V. The membrane was stained with Ponceau S (Cell Signaling catalog #59803) according to the manufacturer’s instructions to confirm protein transfer. The membrane was washed in TBST (VWR catalog #10791-794) with added 0.05% Tween 20 (final 0.1% Tween 20) and blocked in TBST with 5% (w/v) nonfat milk (Amazon, 138 Foods) for an hour, then incubated at 4 °C overnight in TBST with 5% (w/v) BSA and anti-HLA-G (Cell Signaling catalog #79769, 1:1000) and α/β-tubulin (Cell Signaling catalog #2148, 1:1000). The membrane was washed with TBST and incubated in TBST + 5% (w/v) nonfat milk and anti-rabbit IgG HRP-linked antibody (Cell Signaling catalog #7074, 1:1000) for 1 hour then washed in TBST. Bio-Rad Clarity Western ECL Substrate (catalog #1705061) was used according to the manufacturer’s instructions for a 5-minute incubation after a deionized (DI) water rinse. The membrane was imaged with Analytik Jena^TM^ UVP ChemStudio with an optimized exposure time using VisionWorks software.

### Cell Viability

Cells were stained with 2 µM CalceinAM (Invitrogen catalog #C1430) and 1 µM ethidium homodimer (Invitrogen catalog #E1169) for 30 minutes at 37 °C on day 6 to assess cell viability in 3D hydrogels using confocal imaging on a Leica SP8 White Light Confocal. Max-projected z-stack images were analyzed using FIJI.^31^ Average live cell size is presented as the average size of particles in the CalceinAM channel, and percent viability was calculated using the total area of the CalceinAM channel over the total area of the CalceinAM and ethidium homodimer combined. The same threshold, size, and circularity constraints were used for all images. Three Matrigel image values for average live size were disregarded because the cell clusters were indistinguishable, which led to the whole image area being measured as a singular area.

### Cell Metabolism

The metabolic activity of the cells in various culture conditions was assessed using AlamarBlue (Invitrogen catalog #DAL1100) according to the manufacturer’s instructions for 90 minutes at 37 °C using triplicate technical replicates per well on days 0, 2, 4, and 6. Fluorescent values were read using the BioTek Synergy H1 Plate Reader using a gain of 50 at ex/em 560/590. Cell-specific media was used as a background control and subtracted from each sample. Normalized values were obtained by dividing each hydrogel’s metabolic activity by the same hydrogel’s day 0 value to account for any differences between cell seeding density that could occur between experiments.

### Proteomics

CT cells were encapsulated in degradable synthetic (PEG-VPM-GFOGER) hydrogels, degradable natural (Matrigel) hydrogels, or 2D-cultured for 6 days with media changes on days 2 and 4 as detailed above. On day 6, 2D cells were washed with DPBS and lifted with TrypLE. Hydrogels were washed in DPBS for 10 minutes to remove serum. Cell pellets and hydrogels were snap-frozen in liquid nitrogen for lysis and stored at −80 °C until sample preparation and analysis.

Samples were lysed in 8M urea lysis buffer using tip sonication (3 bursts of 30 seconds each with 50% amplitude and a 1-minute interval between bursts). Samples were then spun at 16,000 x gravity for 10 minutes at 4 °C. Protein concentration of clarified lysate was determined using the BCA Protein Assay (Pierce). Protein (25 µg per sample) was digested overnight at 37 °C with trypsin endoproteinase (Trypsin Gold, Promega), and the digested peptides were desalted using 1 cc C18 cartridges (Waters). The resulting tryptic peptides were quantified by BCA assay, and 500 ng of peptides per sample was injected into the column for Data Independent Acquisition (DIA) - Mass Spectrometry (MS) analysis. To generate a spectral library for DIA analysis, equal amounts of peptides per sample were combined to create a pooled library sample fractionated by high pH Reverse Phase Chromatography (HPLC). Pooled peptides (270 µg) were separated on a 25 cm C18 column (Waters XBridge Peptide BEH C18, 130 Å pore size, 3.5 µM particle size, 4.6 mm ID) using a 96-minute ternary gradient formed by liquid chromatography (LC)-MS grade water, LC-MS grade acetonitrile, and 50 mM ammonium hydroxide in LC-MS grade water. Fractions were collected every minute on a 96-deep-well plate and later combined into the final 12 fractions for MS analysis (Pubmed ID: 35977718). MS data was acquired on an Ultimate U3000 RSLCnano LC system coupled to a Thermo Orbitrap Eclipse mass spectrometer equipped with a FAIMS Pro interface (ThermoFisher Scientific). Library fractions were acquired in Data-Dependent Acquisition (DDA) mode, and the individual study samples were acquired in DIA mode. Samples or library fractions in 5 µL injection volume were directly loaded on a C18 column (EASY-Spray ES903 50 cm) and separated by a 2-hr gradient formed by solvent A (LC-MS grade water with 0.1% formic acid) and solvent B (LC-MS grade acetonitrile with 0.1% formic acid). Library runs were acquired in DDA mode with the following settings: cycle time of 3 seconds, FAIMS CV at −45, MS1 scans in the Orbitrap at 120K resolution, a mass range of 380-1480 m/z, and MS2 scans from HCD fragmentation (33% NCE) of the most abundant precursors from MS1 in the Orbitrap at 30K. Samples were acquired in DIA mode with the following settings: cycle time of 3 seconds, FAIMS CV at −45, full scan over a mass range of 380-1080 m/z in the Orbitrap at a resolution of 60K, DIA scans from HCD fragmentation (NCE 33%) across 44 16 m/z wide isolation windows in the Orbitrap at 30K resolution, and a window overlap of 1 m/z.

Spectral Libraries were generated from the 12 library fractions by the Pulsar search engine in Spectronaut using default settings. MS data was searched in Spectronaut against generated spectral libraries with cross-run normalization turned “off,” trypsin-specific digestion, and peptide and protein false discovery rate (FDR) set to 1% as search parameters. Peptide modifications were set for carbamidomethyl (C) and oxidation (M). Resulting protein abundances were normalized using variance stabilizing normalization in R (v4.1.2). Heatmaps and Principal Component Analysis (PCA) plots were created in R (v4.4.0). Heatmaps were generated after log2-transforming the normalized data, employing hierarchical clustering of samples and proteins based on Euclidean distance. For PCA plots, missing values from the log2-transformed normalized data were imputed with zeros.

Contamination by the PEG hydrogel and Xeno (mouse) proteins from media components and Matrigel was evaluated by comparing protein abundances of cells only, cells in hydrogels, and hydrogels only and determined not to impact the protein identifications.

### Enzyme-linked Immunosorbent Assay (ELISA)

Cell supernatants were collected on day 6, centrifuged to remove cell debris, and stored at −80 °C until analysis. Human chromogranin beta (hCGβ) secretion from the cells was determined by ELISA (R&D Systems; catalog #DY9034) with the corresponding ancillary reagent kit according to the manufacturer’s instructions using duplicate technical replicates per sample. Values were read using the BioTek Synergy H1 Plate Reader. A four-parameter logistic curve was used to determine concentrations using MyAssays.com.

### Statistics and Analysis

Image analysis, metabolic activity, and protein secretion were graphed and analyzed using GraphPad Prism 10. Ordinary one- or two-way ANOVAs with Dunnett’s or Tukey’s multiple comparisons tests, as indicated in the figure caption, were used for statistical significance. Data are shown as individual hydrogel replicates with mean and SEM unless otherwise stated.

## RESULTS

### Tunable synthetic hydrogels support TSC-derived placental organoid growth and differentiation comparably to gold standard Matrigel

Recent pioneering studies^8,9,32–36^ have generated placental organoids from TSCs, using a hydrogel matrix to recapitulate a 3D structure (Figure 1A). Traditionally, organoid researchers use the commercially available ECM- and growth factor-rich matrix Matrigel for organoid generation. Matrigel and its growth factor-depleted competitor, Geltrex, are derived from mouse sarcoma, and their natural sourcing contributes to batch-to-batch differences in composition, which may limit experimental reproducibility.^10^ To evaluate whether Matrigel lot-to-lot composition differences could lead to variability in TSC organoid differentiation, we cultured TSCs in two separate lots of Matrigel for 6 days in CT, ST, or EVT differentiation media. We observed distinct differences in HLA-G expression, which is typically highest in the EVT phenotype, in ST condition-exposed TSCs cultured in two different Matrigel lots (Figure 1B, S1A-C, Supporting Information). Standard 2D cultured CT differentiated into ST and EVT subtypes by the same protocol demonstrate trophoblast characteristic markers by flow cytometry (Figure S1D, Supporting Information).

**Figure 1.**
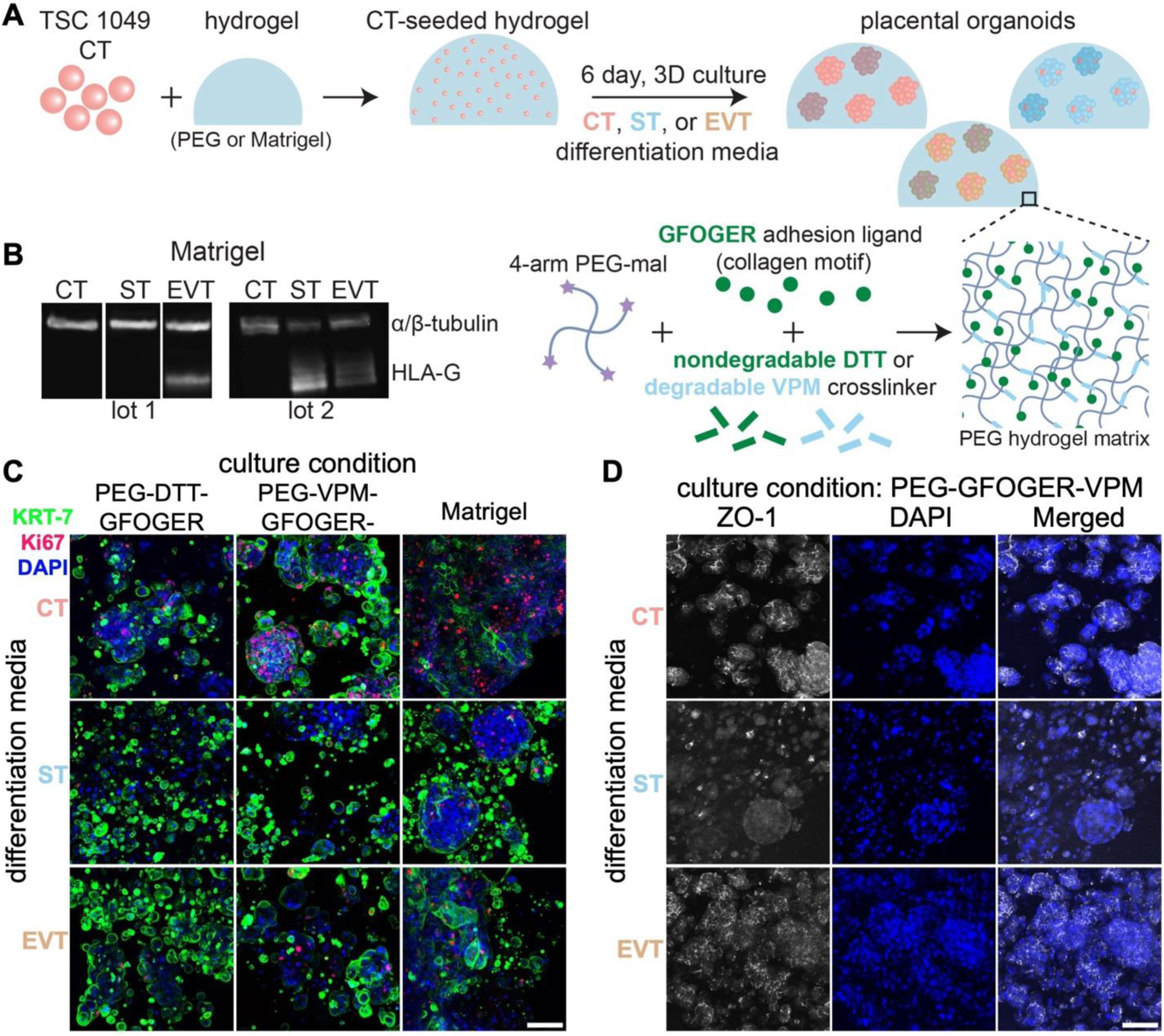
Synthetic hydrogels support trophoblast stem cell (TSC)-derived placental organoid generation comparably to gold standard Matrigel. (A) Schematic of cytotrophoblast (CT) encapsulation into 4-arm poly(ethylene glycol)-maleimide (PEG-mal) modified with collagen-derived ECM motif GFOGER and crosslinked with either protease-degradable VPM or nondegradable DTT and cultured in CT, ST, or EVT differentiation media for 6 days. (B) HLA-G trophoblast marker expression from CT, ST, and EVT cultured in two different lots of Matrigel for 6 days and analyzed via western blotting. Only the Matrigel conditions are shown on the lot 1 blot; the entire blot is shown in Figure S1A, Supporting Information. (C-D) Trophoblast marker expression on day 6 of culture with immunocytochemistry staining of (C) pan-trophoblast marker cytokeratin 7 (KRT7) and proliferation marker Ki67 with DAPI in PEG-GFOGER-DTT, PEG-GFOGER-VPM, and Matrigel, and (D) tight junction marker ZO-1 and DAPI in PEG-GFOGER-DTT. Scale bar = 200 µm. See also Figure S1-2, Supporting Information.

Synthetic ECM-mimicking hydrogels are increasingly used as alternatives to natural matrices in organoid culture due to their controllable, defined properties and limited batch-to-batch variability, which could enhance experimental reproducibility.^10^ We next evaluated whether a PEG-based ECM-mimicking synthetic hydrogel system could support TSC-derived placental organoid growth and differentiation comparably to Matrigel (Figure 1C, S2A, Supporting Information) after 6 days of exposure to CT, ST, or EVT differentiation media. Overall morphology was comparable between Matrigel control cultures and degradable (VPM) or nondegradable (DTT) crosslinked PEG hydrogels, with consistent expression of the pan-trophoblast marker cytokeratin 7 (KRT7) in all groups. CTs retained the highest expression of proliferation marker Ki67. Further, staining for tight-junction protein ZO-1 demonstrated syncytialization via loss of ZO-1 expression in PEG-cultured ST (Figure 1D), confirming that PEG hydrogels can support phenotype-specific differentiation.

### Synthetic hydrogels support placental organoid growth, viability, and function regardless of protease degradability

Synthetic hydrogels enable unique control over the organoid 3D culture environment, such as tunable matrix crosslinkers that degrade in response to specific matrix metalloproteinases (MMP) or nondegradable crosslinkers. As such, we next sought to investigate the influence of synthetic hydrogel crosslinker composition on trophoblast survival, metabolic activity, and functional secretion (**Figure 2**, S3, Supporting Information). We hypothesized that degradable synthetic hydrogels would produce placental organoids with comparable behavior to gold standard Matrigel, particularly for the migratory EVT phenotype. We cultured TSCs in a library of synthetic hydrogels containing collagen-derived GFOGER adhesive ligand and crosslinked with MMP-sensitive peptides (VPM, GDQ, and GPQ-W) and/or nondegradable linear crosslinkers of varying length (DTT: 154 Da, PEG-DT: 1000 or 3400 Da). We included nondegradable natural matrices alginate and agarose, which do not present adhesive ligands to encapsulated cells, and degradable ECM matrices Matrigel and collagen IV, as well as traditional 2D culture, as controls (Figure 2A).

**Figure 2.**
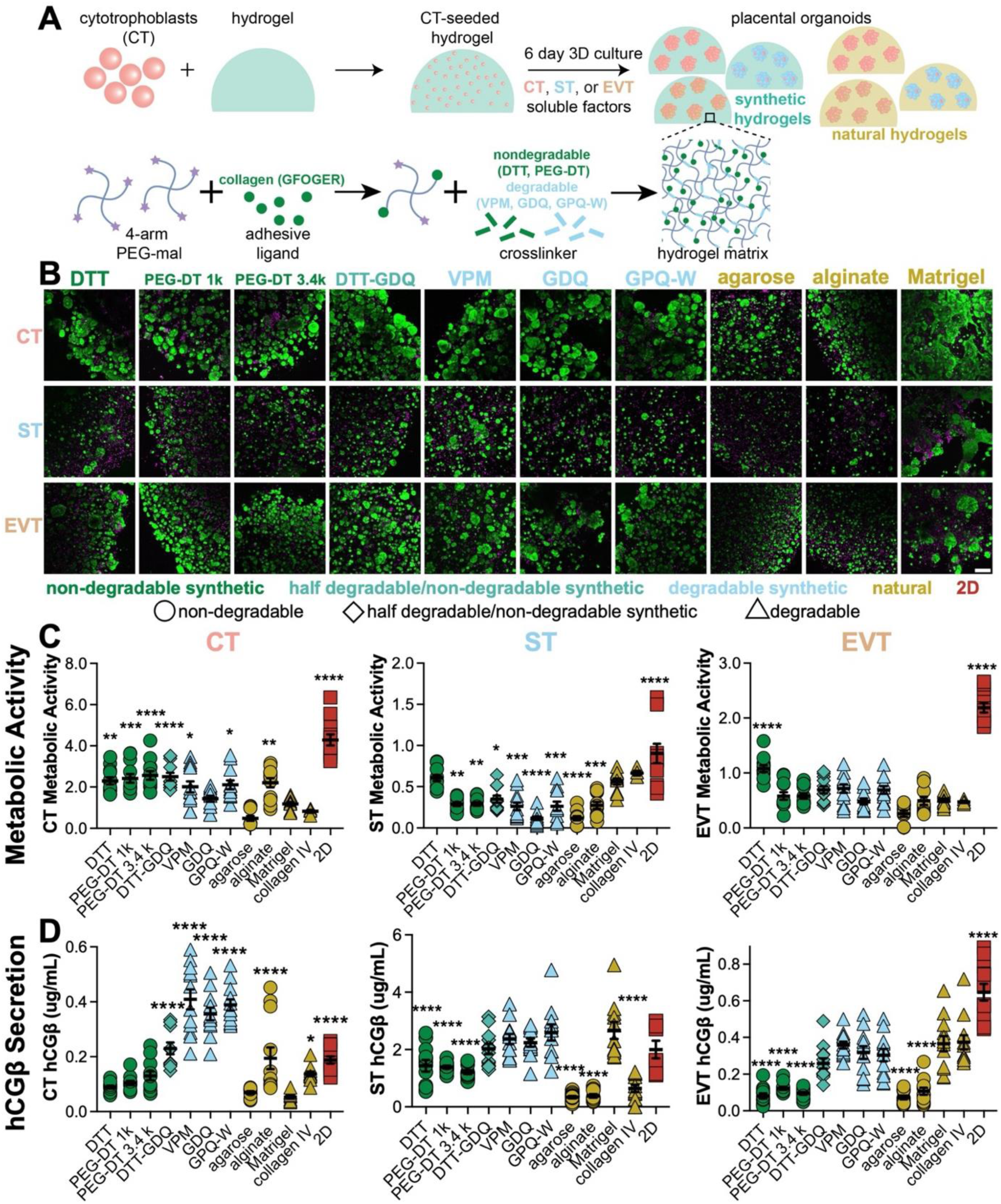
Degradable and nondegradable synthetic hydrogel matrices support placental organoid viability and function comparably to gold standard Matrigel. (A) Schematic of CT encapsulation into PEG-mal functionalized with collagen-derived adhesion ligand GFOGER and crosslinked with protease degradable (VPM, GDQ, or GPQ-W), nondegradable (DTT, PEG-DT), or half nondegradable/half degradable (DTT-GDQ) crosslinkers then cultured for 6 days in CT, ST, or EVT differentiation media and compared to encapsulation in natural degradable (Matrigel, collagen IV), natural nondegradable and nonadhesive (agarose, alginate), or 2D culture. (B) Representative confocal max-projected images of cell viability on day 6 (live – green, dead – magenta) of CT, ST, and EVT phenotypes. (C) Cell metabolic activity via alamarBlue assay on day 6 for CT, ST, and EVT, normalized to day 0. (D) Secretion of hCGβ on day 6 in CT or chemically differentiated ST or EVT phenotypes. n=12 from 3 independent experiments. Data are shown as mean ± SEM and analyzed by ordinary one-way ANOVA with Dunnett’s multiple comparisons test compared to Matrigel; * p < 0.05, ** p < 0.01, *** p < 0.001, **** p < 0.0001. Scale bar = 200 µm. See also Figure S3, Supporting Information.

After 6 days of culture in each hydrogel under CT, ST, or EVT differentiation media conditions, we evaluated organoid viability and morphology via live/dead imaging (Figure 2B, S3A-D, Supporting Information) and metabolic activity via alamarBlue (Figure 2C, S3E-F, Supporting Information). CT viability, organoid average area, and diameter in synthetic hydrogels were comparable to gold standard Matrigel regardless of degradability, and Matrigel organoids were the least circular of all groups (Figure S3A-D, Supporting Information). The metabolic activity of CT organoids cultured in synthetic hydrogel matrices was higher than in Matrigel (Figure 2C). STs are an inherently unstable trophoblast phenotype,^37–40^ and metabolic activity declined across all groups after 2-4 days in culture (Figure S3F, Supporting Information). However, Matrigel supported the highest ST viability, largest organoid area, largest organoid diameter, and reduced circularity compared to all other culture groups (Figure S3A-D, Supporting Information). Finally, EVT media-cultured organoids exhibited significantly higher viability in synthetic hydrogels than Matrigel but comparable organoid area, diameter, circularity, and metabolic activity. Interestingly, nonadhesive and nondegradable alginate and agarose hydrogel controls exhibited consistently reduced viability and organoid size across all trophoblast subtype culture conditions (Figure S3A-E, Supporting Information). This data suggests that both degradable and nondegradable synthetic hydrogels can support placental organoid growth and viability comparably to Matrigel across all differentiation conditions and that local ECM cues are an essential driver of organoid viability in 3D culture.

Human chorionic gonadotropin beta (hCGβ), a pregnancy hormone primarily secreted by the ST phenotype, was also significantly influenced by TSC culture condition (Figure 2D). As expected, secretion of hCGβ was 10-fold higher in the ST phenotype relative to CT and EVT phenotypes. CT and EVT cultured in degradable PEG hydrogels had roughly two to three times higher secretion versus nondegradable PEG hydrogels, suggesting degradable synthetic hydrogels may skew cells to a more ST-like phenotype. Interestingly, natural and synthetic nondegradable hydrogels had significantly lower hCGβ secretion than degradable gels in the ST and EVT phenotype, suggesting that 3D hydrogel degradability may have a greater influence over trophoblast phenotypes than ECM cues. Further, degradable hydrogel hCGβ secretion was comparable to 2D cultured trophoblast despite lower metabolic activity. This suggests that although trophoblast growth is constrained within a 3D environment, phenotypic function is enhanced relative to traditional 2D culture. Overall, these data suggest that 3D placental organoid systems may produce more functional placental-like cultures than 2D culture and that synthetic hydrogel systems can replicate standard Matrigel systems with potentially reduced inter-experiment variability.

### Integrin binding in synthetic hydrogels modulates placental organoid phenotype and function

In our previous study,^22^ trophoblast-like choriocarcinoma cell lines favored degradable synthetic hydrogels; however, in our current study, TSCs grown in nondegradable synthetic hydrogels exhibited comparable metabolic activity and viability to Matrigel (Figure 2). Further, functional differences between placental organoids cultured in collagen, Matrigel, and natural matrices without adhesive ligand presentation (Figure 2) led us to hypothesize that synthetic hydrogel adhesive ligand presentation could alter placental organoid phenotype and function. As such, we selected the nondegradable DTT-crosslinked PEG matrix to evaluate the influence of synthetic hydrogel adhesive ligand presentation on TSC-derived placental organoid behavior (**Figure 3**, S4, Supporting Information). We modified the adhesive ligand in DTT-crosslinked PEG hydrogels to display placenta-expressed ECM motifs (Figure 3A), such as collagen-derived GFOGER, fibronectin-derived RGD or its nonadhesive control RDG, and laminin-derived YIGSR or IKVAV. Confocal viability imaging on day 6 (Figure 3B) and subsequent image analysis (Figure S4A-D, Supporting Information) revealed comparable percent viability of CT and EVT cultured in synthetic PEG with varying adhesion ligands compared to Matrigel. In contrast, the viability of ST cultured in synthetic PEG was lower compared to Matrigel (Figure S4A, Supporting Information), as observed in our previous experiment (Figure 2). CT, ST, and EVT average organoid size were smaller in PEG compared to Matrigel; however, RGD and GFOGER motifs resulted in CT area and CT and EVT diameter comparable to Matrigel (Figure S4B-C, Supporting Information). CT and ST circularity was slightly higher, and EVT circularity was slightly lower, in PEG with all adhesion ligands versus Matrigel. CT organoids grown in hydrogels with RGD and GFOGER motifs exhibited lower circularity more comparable to Matrigel than other motifs (Figure S4D, Supporting Information).

**Figure 3.**
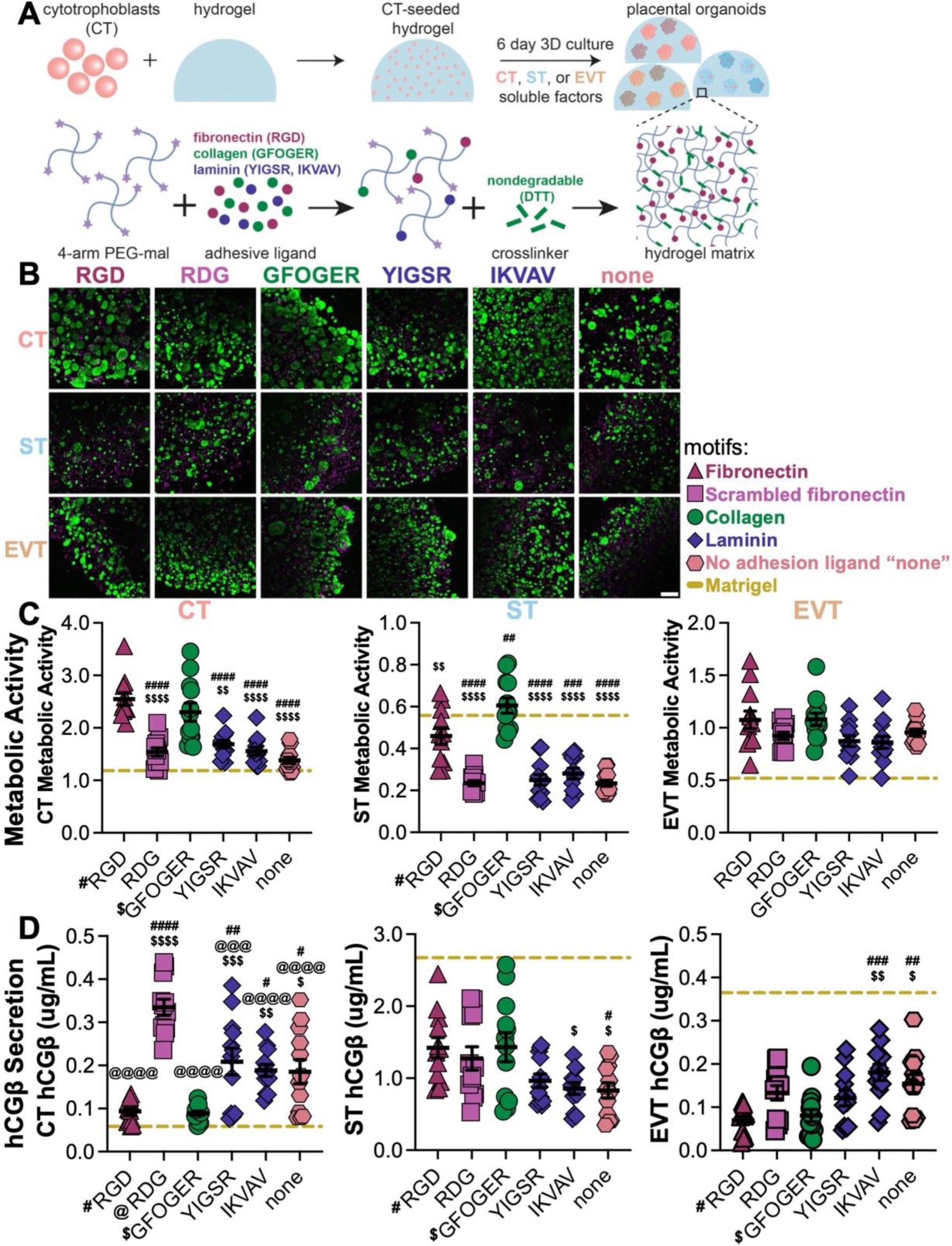
Altering adhesion ligand presentation in nondegradable synthetic hydrogels influences the metabolic activity and function of CT, ST, and EVT-differentiated placental organoids. (A) Schematic of CT encapsulation into PEG-mal functionalized with ECM-derived adhesion ligands (RGD, RDG, GFOGER, YIGSR, IKVAV) or no adhesive ligand (“none”) and crosslinked with nondegradable (DTT) crosslinker then cultured for 6 days in CT, ST, or EVT differentiation media. (B) Representative confocal max-projected images of cell viability on day 6 (live – green, dead – magenta) of CT, ST, and EVT phenotypes in synthetic PEG hydrogels using DTT nondegradable crosslinker and varying adhesion ligands. (C) Cell metabolic activity via alamarBlue assay on day 6 for CT, ST, and EVT, normalized to day 0, compared to Matrigel (dashed line). (D) Secretion of hCGβ on day 6 in CT or chemically differentiated ST or EVT phenotypes, compared to Matrigel (dashed line). n=12 from 3 independent experiments. Data are shown as mean ± SEM and analyzed by ordinary one-way ANOVA with Tukey’s multiple comparisons test. # compared to RGD, @ compared to RDG, and $ compared to GFOGER: * p < 0.05, ** p < 0.01, *** p < 0.001, **** p < 0.0001. Scale bar = 200 µm. See also Figure S4, Supporting Information.

The metabolic activity of CT, ST, and EVT was consistently higher in synthetic hydrogels with RGD and GFOGER adhesive ligands (Figure 3C), though not statistically significant in the EVT phenotype. As in our previous experiment (Figure 2C), CT and EVT organoids grown in PEG-DTT hydrogels with varying adhesion ligands had a higher average metabolic activity compared to Matrigel, whereas only ST grown in PEG-DTT hydrogels with GFOGER adhesion ligand had a higher average metabolic activity than organoids grown in Matrigel (Figure 3C). EVT organoid viability, metabolic activity, and hCGβ secretion demonstrated minimal dependence on adhesive ligands, with RGD and GFOGER groups demonstrating slightly lower hCGβ than IKVAV and no adhesion ligand groups (“none,” Figure 3D). As we observed in Figure 2D, the ST organoid hCGβ secretion range was approximately ten-fold that of CT and EVT organoids (Figure 3D). Overall, adhesive ligand inclusion within synthetic hydrogel matrices significantly influenced trophoblast metabolic activity and function, with fibronectin and collagen-derived sequences resulting in placental organoid phenotypic behavior most aligned with expected differentiation condition relative to laminin-derived or control conditions (no adhesive ligand “none” or scrambled adhesive ligand).

### Placental organoid culture in Matrigel encourages an EVT-dominant trophoblast phenotype

TSC metabolic activity, viability, and hCGβ secretion provided insight into the influence of culture condition on trophoblast behavior but provided limited detail of placental organoid phenotype. To evaluate the influence of 2D or 3D culture environments on TSC-derived placental organoid differentiation more comprehensively, we performed liquid-chromatography tandem mass spectrometry (LC-MS) and proteomic analysis on day 6 organoids cultured in CT, ST, and EVT differentiation media (**Figure 4**-7, S5-8, Supporting Information). We compared organoids grown in 3D natural Matrigel and synthetic PEG-VPM-GFOGER hydrogels to standard 2D culture (Figure 4A). Principle component analysis from unsupervised clustering of proteins present in >50% of samples (n=6004 proteins) demonstrates distinct protein expression profiles with predominant clustering of samples by trophoblast differentiation media (Figure 4B) rather than by 2D or 3D culture condition (Figure 4C). Overall, protein expression of CT and EVT grown in 2D and Matrigel conditions cluster closer than when grown in PEG; however, ST grown in PEG and Matrigel cluster closer than when they are grown in 2D. Notably, TSC cultured in Matrigel under ST differentiation shifted closer to the EVT cluster, and TSC in PEG hydrogels under CT differentiation shifted closer to the ST cluster (Figure 4C, right).

**Figure 4.**
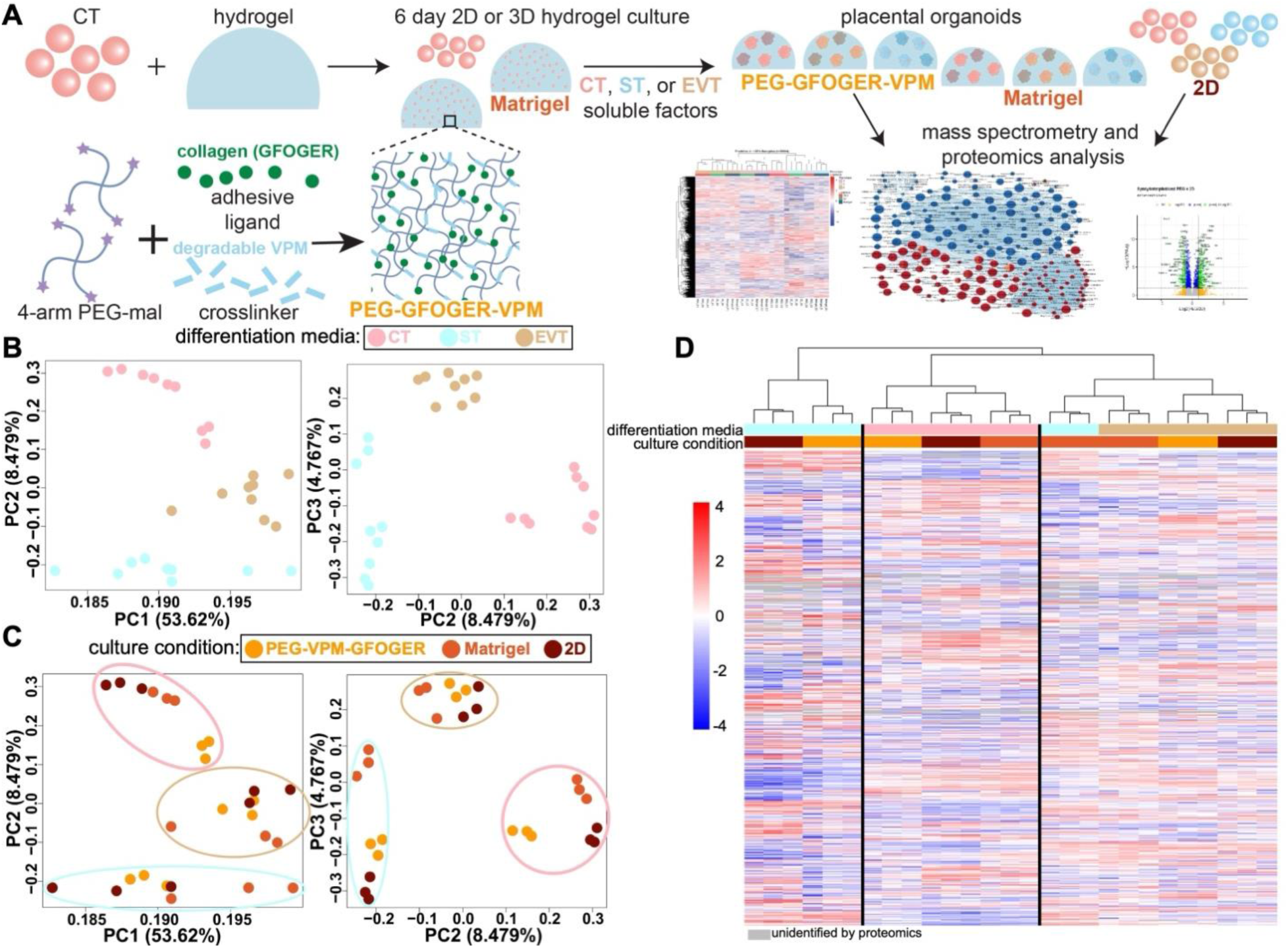
TSC proteomic analysis reveals differentiation media- and culture condition-dependent differential protein abundances. (A) Schematic of CT encapsulation in synthetic degradable (PEG-GFOGER-VPM) or natural degradable (Matrigel) matrices or 2D cultured for 6 days in CT, ST, or EVT differentiation media before using high-throughput mass spectrometry for proteomic analysis. (B-C) Principal component analysis of global protein abundance shows more clustering of (B) CT, ST, and EVT by differentiation media condition than by (C) culture condition. (D) Hierarchical clustering of differentially abundant proteins (p < 0.05) revealed clustering primarily by differentiation condition. See also Figure S5-S6, Supporting Information.

**Figure 5.**
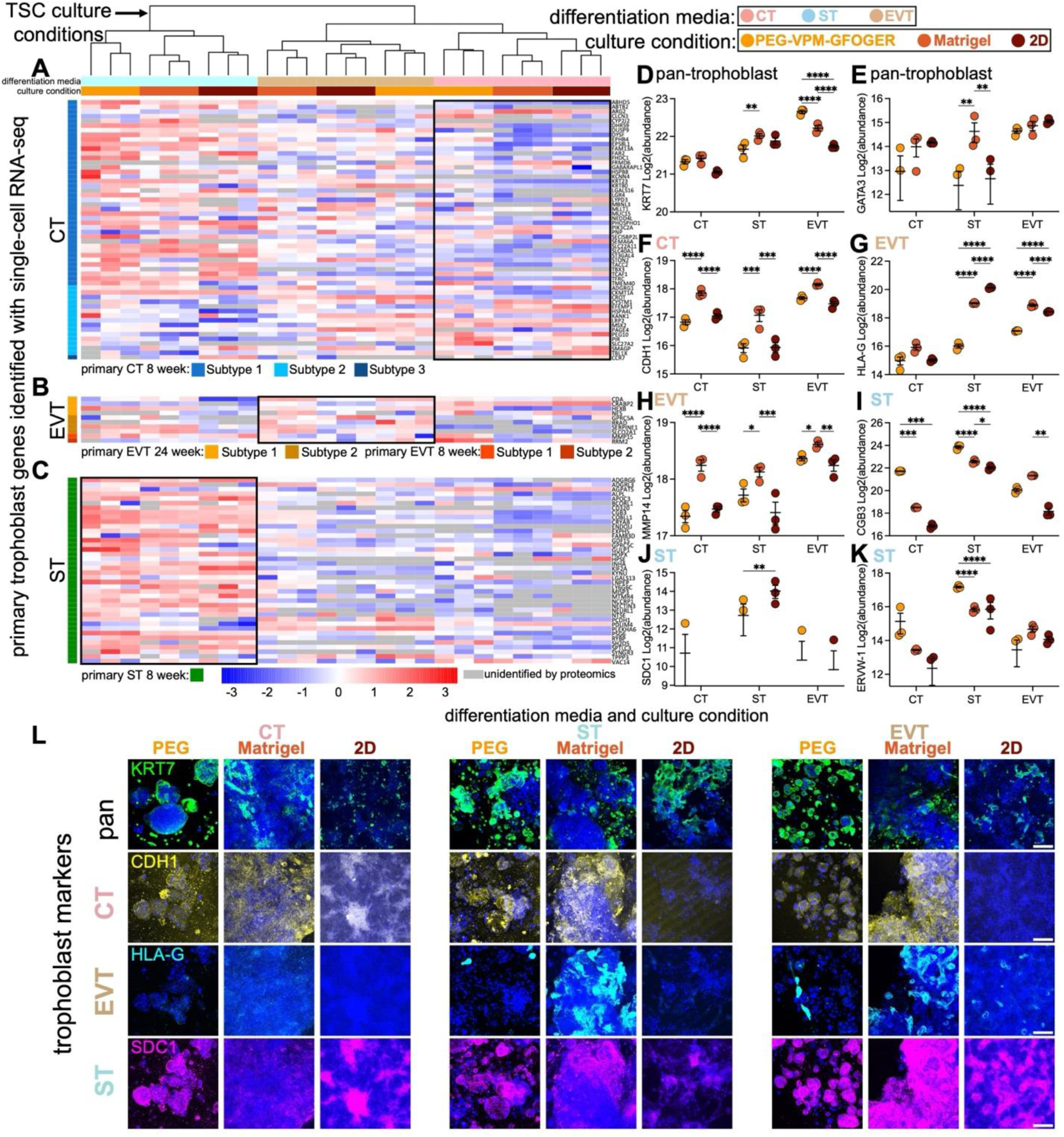
Synthetic hydrogels support ST-like placental organoid phenotypes, while Matrigel skews organoids to an EVT-like phenotype. (A-C) TSC protein abundance markers that align with (A) CT, (B) EVT, and (C) ST 8- or 24-week gestational age primary human trophoblast subtypes identified with single-cell RNA-seq^41^ after culture in PEG-GFOGER-VPM, Matrigel, or 2D for 6 days. Abundances of pan-trophoblast markers (D) KRT7 and (E) GATA3, CT marker (F) CDH1, EVT markers (G) HLA-G and (H) MMP14, and ST markers (I) CGB3, (J) SDC1 and (K) ERVW-1. n=3. Data are shown as mean ± SEM and analyzed by two-way ANOVA with Tukey’s multiple comparisons test: * p < 0.05, ** p < 0.01, *** p < 0.001, **** p < 0.0001. (L) Protein expression via immunocytochemistry of KRT7, CDH1, HLA-G, and SDC1 with DAPI (blue). Scale bar = 200 µm. See also Figure S7, Supporting Information.

**Figure 6.**
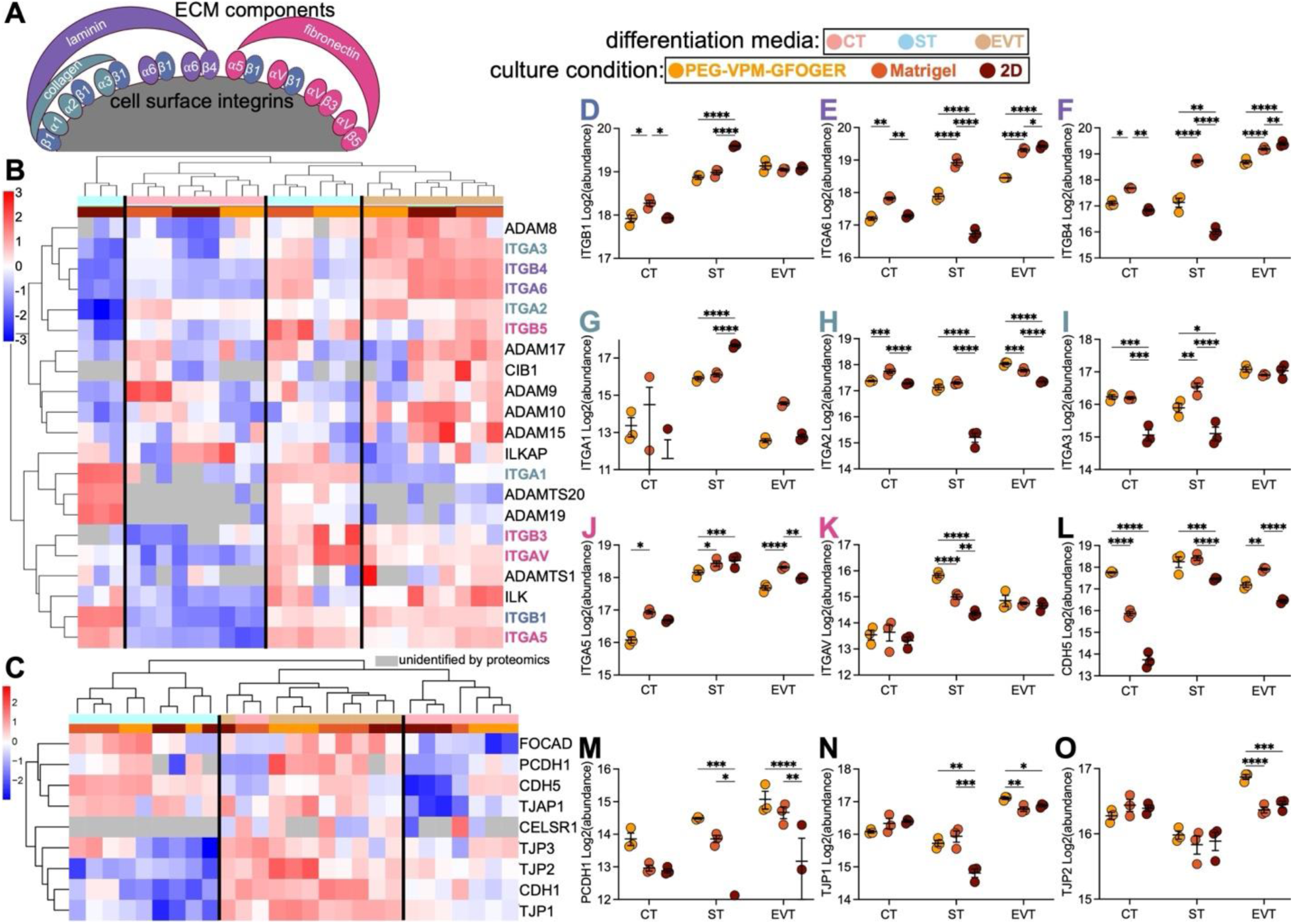
The EVT phenotype expresses the most abundant adhesome proteins. (A) Schematic of integrin complexes for ECM adhesion. (B) Normalized protein abundance of integrins (ITG), a disintegrin and a metalloproteinase (ADAM) family members, and kinase proteins from CT, ST, and EVT cultured in PEG-GFOGER-VPM, Matrigel, or 2D for 6 days. Overview of (B) integrin (ITG), (C) cadherin (CDH), and tight junction protein (TJP) abundances. Normalized protein abundances for (D) fibronectin-, collagen-, and laminin-binding protein ITGB1, (E-F) laminin-binding proteins (E) ITGA6 and (F) ITGB4, (G-I) collagen- and laminin-binding proteins (G) ITGA1, (H) ITGA2, and (I) ITGA3, (J-K) fibronectin-binding proteins (J) ITGA5 and (K) ITGAV, (L) CDH5, (M) PDCH1, (N) TJP1, and (O) TJP2. n=3. Data are shown as mean ± SEM and analyzed by two-way ANOVA with Tukey’s multiple comparisons test: * p < 0.05, ** p < 0.01, *** p < 0.001, **** p < 0.0001. See also Figure S7, Supporting Information.

**Figure 7.**
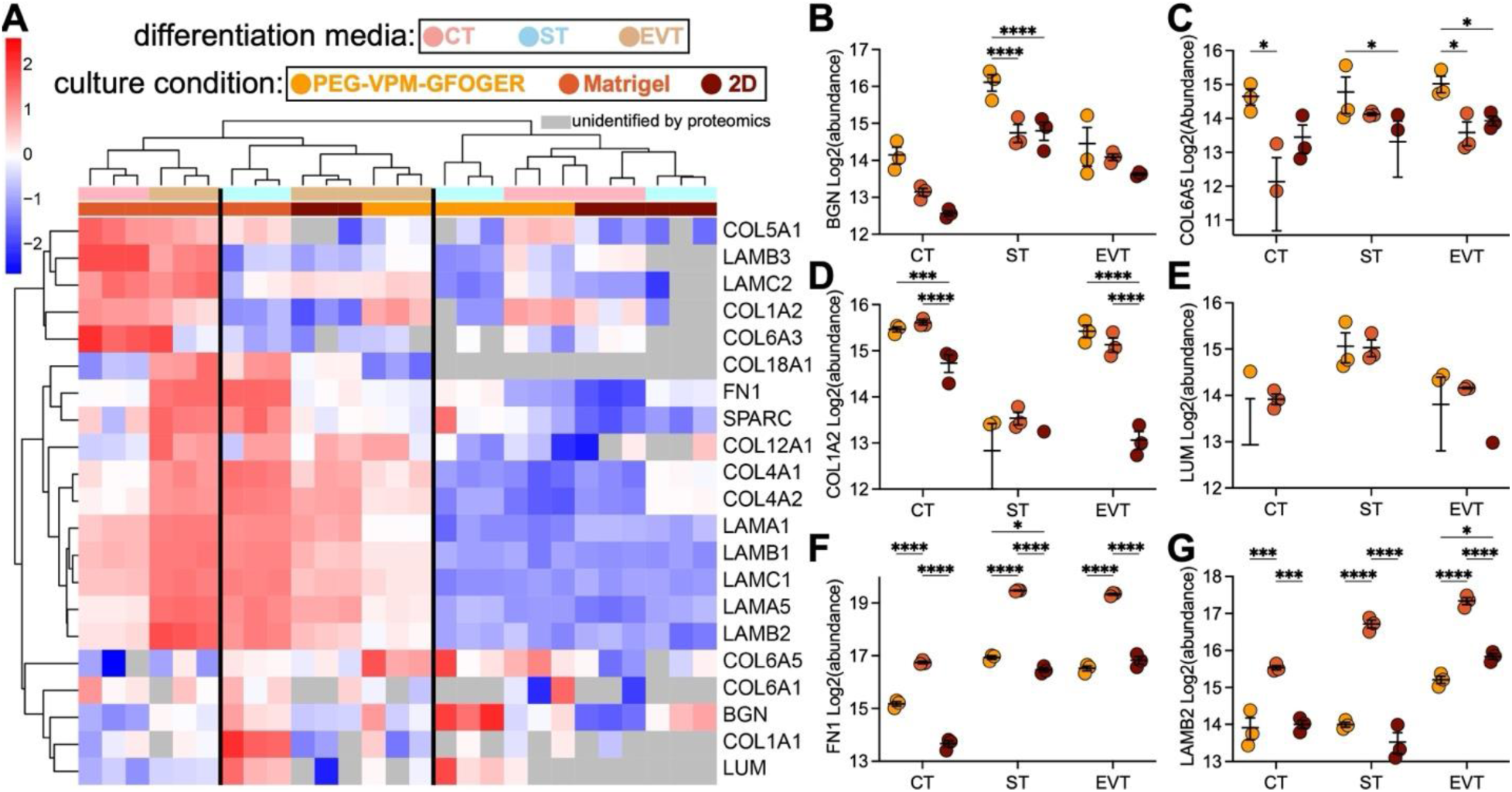
Extracellular matrix (ECM) component abundances are highest in ST and EVT cultured in Matrigel. (A) Normalized ECM abundances from CT, ST, and EVT cultured in PEG-GFOGER-VPM, Matrigel, or 2D for 6 days, then analyzed by mass-spectrometry proteomics. (B-G) ECM abundances of (B) biglycan (BGN), (C) collagen alpha-1(VI) (COL6A5), (D) collagen alpha-1(XXI) (COL1A2), (E) lumican (LUM), (F) fibronectin (FN1), and (G) laminin subunit beta-2 (LAMB2). n=3. Data are shown as mean ± SEM and analyzed by two-way ANOVA with Tukey’s multiple comparisons test: * p < 0.05, ** p < 0.01, *** p < 0.001, **** p < 0.0001.

Between samples, 2968 proteins were differentially abundant (p < 0.05, Figure 4D). Hierarchical clustering of these proteins reveals clustering mainly by cell phenotype; however, Matrigel cultured STs and EVTs clustered together with the 2D and PEG cultured EVT cluster. These data suggest that the soluble differentiation cues present in each condition have the most substantial impact on trophoblast phenotype, while the culture format (e.g., 2D or 3D) can also significantly influence cell phenotype.

Next, we investigated pathway enrichment to identify changes in protein expression across culture conditions (Figure S5, Supporting Information). Overall, cells grown in 3D (PEG and Matrigel) have a higher abundance of proteins associated with cell motility and migration, cytoskeleton organization, and responses to stimulus. In comparison, 2D cultured cells have a higher abundance of proteins associated with metabolic processes and RNA processing pathways. These data are consistent with the increased metabolic activity of 2D cultured cells compared to 3D conditions (Figure 2C). Additionally, we did not see an overall trend in increased apoptosis-associated proteins driven by a specific culture or differentiation media condition (Figure S6D, Supporting Information).

We next sought to evaluate how our differentiated placental organoid phenotypes aligned with primary trophoblast phenotypes (**Figure 5**). First, we broadly evaluated our proteomic protein abundances against genes identified in a single-cell RNA-seq dataset^41^ of primary trophoblast phenotypes (CT, ST, and EVT) isolated at 8- and 24-week gestational ages (n = 1643, Figure 5A-C). TSCs cultured in CT media and 2D, 3D Matrigel, or 3D PEG hydrogels demonstrated greater abundances of proteins from CT subtype 2 of an 8-week gestational age placenta than any other subtype (Figure 5A); however, PEG-cultured CT also showed a higher abundance of subtype 1-associated genes than other culture conditions, a CT subtype identified as a potential ST progenitor.^41^ TSCs cultured in PEG or Matrigel in EVT media had a higher abundance of primary EVT proteins from 24-week gestational age compared to 2D cultured cells in EVT media. These data support our hypothesis that 3D culture skews TSCs toward an EVT phenotype (Figure 5B). TSCs in ST media and cultured in PEG, Matrigel, and 2D all shared abundant proteins with 8-week gestational age STs (Figure 5C). Interestingly, ST-cultured cells also highly expressed CT subtype-1-associated genes, further supporting the association of this subtype with an ST progenitor.

Next, we evaluated specific markers commonly used to identify trophoblast phenotypes in our proteomics dataset (Figure 5D-K) and by immunocytochemistry (Figure 5L). Pan-trophoblast markers KRT7 and GATA3 broadly increased in expression from CT to ST and EVT media culture conditions, with significantly higher protein abundances in Matrigel relative to PEG for ST and higher KRT7 abundance in EVT when cultured in PEG versus Matrigel and 2D (Figure 5D-E), while staining was the most consistent in PEG-cultured CT, ST, and EVT (Figure 5L). CT marker CDH1 was highly abundant in Matrigel-cultured CT, ST, and EVT relative to PEG and 2D conditions (Figure 5F), which was also observed in staining (Figure 5L). EVT-associated proteins HLA-G and MMP14 were generally higher in EVT differentiation and Matrigel culture conditions (Figure 5G, 5H), except for HLA-G, which was highly expressed in 2D-cultured ST; however, HLA-G staining was only highly observed in Matrigel ST and EVT groups (Figure 5L). ST-associated proteins CGB3, SDC1, and ERVW-1 were most abundant in ST differentiation and PEG culture conditions (Figure 5I-K). Overall, proteome analysis of TSC phenotype indicates that PEG hydrogels resulted in more ST-specific protein expression more than Matrigel and 2D culture, and Matrigel resulted in more EVT-associated protein expression than PEG and 2D culture. Further, proteomics and immunocytochemistry demonstrated heterogeneity of phenotypic protein expression within TSC cultures, with CT, ST, and EVT-associated proteins identified in all culture conditions.

### Culture condition and trophoblast phenotype significantly alter the placental organoid adhesome and extracellular matrix production

Our observation that synthetic hydrogel adhesive ligand sequences may drive differences in TSC phenotypic behavior (Figure 3) led us to hypothesize that CT, ST, and EVT may exhibit significant differences in integrin expression and ECM secretion. To investigate this, we used our proteome dataset to evaluate the influence of differentiation media and 2D or 3D culture conditions on the TSC adhesome and ECM production (**Figure 6**-7, S7, Supporting Information). 2D cells were cultured on collagen-coated tissue culture plastic, and PEG hydrogels presented the GFOGER collagen motif, restricting cell engagement to collagen-specific integrins, whereas Matrigel possesses ubiquitous ECM adhesive ligands. As such, we wondered whether these culture conditions altered TSC integrin upregulation (Figure 6A). Broadly, we observed the most significant upregulation in cell-matrix integrin (Figure 6B) and cell-cell cadherin (Figure 6C) adhesome proteins in the EVT culture condition, followed by the ST and then CT conditions. Interestingly, integrin expression was largely clustered by differentiation conditions, except for the 2D ST group, which was an outlier from the entire dataset.

Proteomic analysis revealed the presence of several integrins (ITG, α1, α2, α3, α5, α6, αV, β1, β3, β4, and β5) associated with fibronectin-, collagen-, and laminin-binding (Figure 6A, 6B). ITGB1, which forms complexes with other integrins to bind to fibronectin, collagen, and laminin, had the highest abundance in ST and EVT differentiation conditions, with 2D-cultured ST having significantly higher expression than PEG and Matrigel ST (Figure 6D). Laminin-associated ITGA6 and ITGB4 had the highest abundances in Matrigel-cultured ST and EVT and 2D-cultured EVT (Figure 6E, 6F). Collagen- and laminin-associated ITGA1 abundances were the highest in the 2D ST condition, while ITGA2, ITGA6, and ITGB4 abundances were the lowest in the 2D ST condition and generally the highest in the EVT differentiation condition (Figure 6G, 6H, 6I). Fibronectin-associated ITGA5 and ITGAV were highest in the ST differentiation condition and the lowest in the CT differentiation condition (Figure 6J, 6K). Overall, integrin protein expressions were the highest in ST and EVT differentiation conditions.

Similar to the pattern of integrin expression, most ADAM (a disintegrin and a metalloproteinase) family members (ADAM8, 9, 10, 17, 15, and 19) and ADAM with thrombospondin motif members (ADAMTS1 and 20), which bind to integrins and have cell adhesive properties (Figure 6B), were upregulated in EVT and ST relative to CT. Cadherin family members, which mediate cell-to-cell adhesion and which we expected to be upregulated in organoids over 2D culture, tended to be significantly upregulated in 3D culture relative to 2D (Figure 6C, 6L, 6M). Similarly, tight junction proteins (TJP1, 2, and 3) are expected to be lower in STs given the syncytialization characteristic of this cell type, which we also broadly observed (Figure 6C, 6N, 6O). Overall, integrins, ADAMs, cadherins, and tight junction proteins were the highest in ST and EVT differentiation conditions.

We previously noted that the EVT phenotype demonstrated less functional dependence on synthetic hydrogel adhesive ligand inclusion (Figure 3, S4, Supporting Information), leading us to hypothesize that the EVT phenotype produced its own ECM in 3D culture. As such, we next evaluated whether culture or differentiation conditions altered ECM production by TSCs (**Figure 7**). Proteomic analysis revealed ECM production was clustered primarily by culture condition, with Matrigel exhibiting the greatest upregulation of ECM components biglycan, collagen, decorin, fibronectin, laminin, lumican, osteonectin (SPARC), thrombospondin, and vitronectin, particularly in the ST and EVT differentiation conditions (Figure 7A). Biglycan (BGN) and collagen alpha-1(VI) (COL6A5) abundances were highest in PEG culture conditions within each differentiation media condition (Figure 7B, 7C); however, collagen alpha-1(XII) (COL1A2) and lumican (LUM) protein abundances were highest in 3D (PEG and Matrigel) conditions versus 2D (Figure 7D, 7E). Meanwhile, fibronectin (FN1) and laminin subunit beta-2 (LAMB2) abundances were the highest in all phenotypes cultured in Matrigel compared to PEG and 2D culture conditions (Figure 7F, 7G), which is also evident in volcano plot comparisons showing many ECM proteins with higher abundance in Matrigel versus 2D and PEG (Figure S6A-C, Supporting Information). Overall, Matrigel-cultured ST and EVT had the highest abundances of ECM proteins, while EVT cultured in the three culture conditions generally expressed higher collagen and laminin abundances than ST and CT.

## DISCUSSION

TSC-derived placental organoids enable the study of a relatively inaccessible organ, the early gestation human placenta; however, a reliance on natural 3D matrices to generate placental organoids, such as Matrigel, may introduce interexperimental variability^10^ that limits reproducibility and confidence in results. Traditionally, Matrigel has been used to culture placental organoids ^8,9,32–36^ due to its ready commercial availability, ease of use, and the abundance of growth factors and ECM cues that support trophoblast viability; however, in our hands, Matrigel lot-to-lot variability demonstrated placental organoid phenotype variation that could contribute to reduced experimental reproducibility (Figure 1B). As an alternative, we evaluated a synthetic PEG-based hydrogel system that may provide greater control and reproducibility of placental organoid generation (Figure 1C-D). We found that PEG-based hydrogels generated comparable organoids to Matrigel, demonstrating that ST and EVT differentiation exhibits a comparable decrease in Ki67 expression relative to Matrigel (Figure 1C), an expected finding since the ST phenotype is known to have a reduced proliferative capacity and requires constant replenishment by CT.^37–40^ This was further evident when we evaluated PEG, Matrigel, and 2D-cultured TSCs over 6 days, with consistently reduced metabolic activity in ST differentiation media-exposed TSCs (Figure 2C, 3C, S3E-F, S4E-F, Supporting Information); however, we did not see a significant increase in apoptosis-associated proteins in ST conditions compared to CT and EVT conditions (Figure S6D, Supporting Information).

To mimic Matrigel’s permissive environment, cell-responsive degradability was incorporated into synthetic PEG hydrogels via VPM, GDQ, and GPQ-W peptides, which have varying protease sensitivity^22–29^ (Table 1). Proteomics results revealed the production of MMP-2, −12, −14, and −15 in various trophoblast conditions (Table S1, Supporting Information), which can degrade these peptide linkers. Unexpectedly, TSC grown in nondegradable PEG hydrogels had higher metabolic activity and lower hCGβ secretion than protease-degradable PEG (Figure 2C, 2D). Though agarose and alginate are also nonprotease-degradable, metabolic activity and hCGβ secretion were both low and did not show favorable trends. Interestingly, agarose and alginate generally produced lower percent viability and smaller average organoids than Matrigel (Figure S2, Supporting Information). However, alginate metabolic activity and hCGβ secretion were significantly higher than Matrigel for CT organoids (Figure 2C, 2D) while comparable or lower for ST and EVT differentiation conditions. Alginate and agarose both lack growth factors and native adhesive ligands, and this implies that ST and EVT may rely more on cell-matrix cues than CT. However, matrix stiffness, which can regulate hormone production,^42^ may play a role as alginate and agarose (4867 and 5483 Pa, respectively)^43^ have greater stiffnesses more comparable to the decidua basalis (1250 Pa) due to invading EVTs,^44^ whereas PEG (50 Pa)^43,45^ and Matrigel (∼113 Pa, lot-to-lot variability)^44^ have more comparable stiffnesses to the placenta (232 Pa).^44^ Overall, placental organoid function was not solely dependent upon matrix degradation but relied significantly on soluble and matrix environmental cues.

Integrin binding can impact numerous cell processes, including differentiation, proliferation, and gene expression. In our previous studies with choriocarcinoma-like trophoblast cell lines, adhesive ligands did not significantly influence cell function.^22^ By contrast, primary TSCs exhibit phenotype-dependent responsiveness to matrix-bound environmental cues. Collagens I, III, IV, and VI and laminin represent major components of the placenta ECM.^46^ As such, we included collagen (GFOGER) and laminin (YIGSR and IKVAV) ECM-derived adhesion ligands into our PEG hydrogel system, as well as the ubiquitous ligand RGD, identified initially on fibronectin but also present on vitronectin and osteopontin, among others.^47,48^ Interestingly, RGD and collagen ligands better supported TSC metabolic activity and hCGβ secretion than laminin adhesion ligands (Figure 3C, 3D). Similar to the PEG condition when varying degradability (Figure 2), the highest metabolic activity conditions for CT and EVT, RGD and GFOGER adhesive ligands, were the lowest hCGβ-secreting conditions. In contrast, the highest metabolic conditions for ST (RGD and GFOGER) translated to a higher secretion of hCGβ (Figure 3C, 3D). This correlation may be due to the lower metabolic activity of the ST phenotype, which is responsible for secreting the majority of the hCGβ in the placenta.^1,49–51^ As ST cultured with RGD and GFOGER adhesive ligands secreted significantly more hCGβ, while CT and EVT secreted significantly less hCGβ, this suggests RGD and collagen adhesive ligands may assist in stabilizing TSC phenotype. Matrigel-cultured CT and EVT had slightly lower metabolic activity than PEG hydrogels with each adhesion ligand tested, whereas ST cultured with RGD and GFOGER ligands had comparable metabolic activity to Matrigel (Figure 3C), suggesting a higher dependency on fibronectin and collagen ECM for the ST phenotype.

We hypothesized that CT would favor RGD, GFOGER, and IKVAV due to their expression of α5β1, α1β1, and α6β1,^52–60^ respectively, and we saw an increase in the average live size and metabolic activity of CT when cultured with RGD and GFOGER. We also hypothesized that ST would favor RGD due to their expression of α5β1 and αVβ3,^53,56,58^ and while ST organoids exhibit higher metabolic activity with RGD, this also occurred with GFOGER. STs mainly function to provide barrier protection, so they may bind less and present fewer integrins than other cell types due to their syncytialization. Further, we hypothesized that EVT would favor RGD, GFOGER, and IKVAV due to their expression of αVβ3, α1β1, and α6β1,^52–55,57,60–63^ respectively. We saw an increase in EVT’s average size and metabolic activity when cultured with RGD and GFOGER; however, there were no significant differences. PEG hydrogels with all tested adhesion ligands produced higher metabolic activity than Matrigel in CT, ST (GFOGER only), and EVT differentiation conditions. Additionally, we saw evidence of higher ECM abundances, particularly collagen and laminin, from EVT, as we suspected based on their independence from adhesion ligands (Figure 7), which could be attributable to the influence of growth factors provided by the Matrigel or the additive measurement of ECM proteins present in Matrigel.

As the PEG hydrogels included the GFOGER collagen motif, Matrigel contains a high laminin content,^64^ and 2D CT/ST and EVT were cultured on collagen and fibronectin, respectively, we questioned whether we would observe integrin expression associated with these ECM; however, we did not see significant evidence that these culture conditions were inducing integrin changes (Figure 6, S7, Supporting Information). For example, 2D cultured CT and ST and Matrigel cultured phenotypes did not have a significant presence of ITGA1 compared to 2D cultured EVT cultured on fibronectin. These data suggest that including binding proteins may not contribute to increased integrin protein expression alone and is mainly modulated by trophoblast phenotype. This aligns with previous studies suggesting that trophoblasts require direct contact with endothelial cells for the upregulation of ITGB1, and secreted endothelial factors are not enough.^65^ ITGA1 and HLA-G are co-expressed on a population of EVT, while NOTCH1 and ITGA5 are two column trophoblast states,^66^ which means 2D cultured ST may behave more like EVT cells and explains the lower expression of ST markers in this condition (Figure 5, 6, S7, Supporting Information). In the first trimester, ITGB3 expression is localized to ST,^67^ and ST culture in PEG has the highest expression of ITGB3 (Figure 6, S7, Supporting Information). These data are essential characteristics of trophoblasts as integrin expression may determine trophoblast behavior and differentiation.^60^

Proteomics results revealed greater differential protein expression between differentiation media conditions than between 2D or 3D culture conditions, suggesting a greater influence of microenvironment chemical cues versus mechanical cues over trophoblast differentiation (Figure 4B); however, culture conditions also significantly affected phenotypic protein expression (Figure 4C-D, 5-7). Trophoblast markers KRT7, GATA3, and transcription factor AP-2α (TFAP2C) were present in all culture and differentiation media conditions (Figure 5-E, S7A-C, Supporting Information). GATA3 and TFAP2C are considered mononuclear trophoblast markers as reports have shown their absence on ST phenotypes;^68–71^ however, GATA3 and TFAP2C expression were highest in Matrigel and 2D cultured ST, respectively, possibly signaling these methods do not lead to typical ST phenotypes or that this expression varies with gestational age. ST phenotypes have some of the most defined markers^8^ and many of these were most highly upregulated in PEG culture (Figure 5I-K, S7F-K, Supporting Information), supporting our conclusion that PEG better supports the ST phenotype than Matrigel and 2D culture. Conversely, EVT markers HLA-G and MMP14 were high in Matrigel cultured ST and EVT, suggesting Matrigel better supports EVT differentiation than PEG and 2D culture and skews cells to an EVT-like phenotype (Figure 5G-H, S7L-N, Supporting Information).

Previous studies using Matrigel have generated trophoblast organoids using TSC with inverted placenta structures (CT on outside, ST on inside) or physiological-like organization (EVT or ST on outside, CT on inside).^8,9,32–36^ However, in our hands, TSC created a densely population cell sheet that degraded Matrigel within a few days (Figure 2B) with no visible structure modeling (Figure 5L). This is likely due to differing protocols, where cell number and Matrigel concentration play a key role in trophoblast differentiation. Additionally, our CT, ST, and EVT differentiation media slightly differed from previously reported differentiation media. Some studies do not include Noggin and SB431542 or include R-spondin and PGE2,^8,72^ which may significantly affect cell differentiation.

### Limitations of the study

Like other organoid systems, there are limitations to this model and the application of its results. Notably, these results are specific to a single donor TSC, and results may vary with TSC gestational age, donor sex, and culture time. Here we used previously established differentiation media,^6^ which includes 0.5-2% Matrigel for EVT differentiation, so Matrigel batch-to-batch variability could also affect EVT differentiation in synthetic hydrogels. Future studies could assess EVT differentiation in synthetic hydrogels without the addition of Matrigel or apply a more defined combination of ECM and growth factors. This model also used a higher cell number than previous Matrigel cultured models,^8,9,32–36^ which may limit applications where only low trophoblast numbers can be isolated, such as for personalized medicine applications where chronic villus sampling procedures are used.^32^

Furthermore, we did not confirm trophoblast lineage against all four criteria of primary first-trimester trophoblast: (1) methylation of ELF5, (2) mRNA levels of C19MC, (3) protein markers KRT7, GATA3, and TFAP2C, and (4) expression of HLA class I molecules (HLA-A^-^, HLA-B^-^, and HLA-G^-/+^.^68^ Proteomics results revealed the presence of GATA3, KRT7, TFAP2C, HLA-B, HLA-E, and HLA-G (Figure 5D, 5E, 5G, S7, Table S1, Supporting Information). Additionally, we did not investigate the spatial distribution of cells within the organoids in this model through high-resolution imaging; we confirmed the presence of these common markers by western blotting, confocal imaging, and proteomic analysis. Finally, in addition to trophoblasts, the human placenta is comprised of other cells, including fibroblasts, pericytes, and mesenchymal-derived macrophages (Hofbauer cells) that aid in trophoblast differentiation and angiogenesis,^73,74^ whereas this model only investigated trophoblast differentiation through chemical and mechanical signaling. The incorporation of these supporting cells could result in a more faithful representation of placental tissue and behavior in this organoid model.

## CONCLUSION

We have engineered a synthetic PEG-mal hydrogel system with tunable degradability and ECM-derived adhesive ligands capable of supporting the differentiation and function of TSC-derived placental organoids and compared this model to gold standard Matrigel and 2D culture. We found that viability, metabolic activity, protein secretion, and protein expression of placental organoids were significantly affected by differentiation conditions (CT, ST, or EVT) and cell culture conditions (PEG, Matrigel, or 2D). Generally, PEG hydrogel-cultured placental organoids favor an ST phenotype, while Matrigel-cultured organoids favor an EVT-associated phenotype. Overall, we demonstrated that well-defined and tunable PEG hydrogels can generate placental organoids containing CT, ST, and EVT phenotypes, enabling the study of trophoblast cell-cell and cell-matrix cues. This hydrogel system may enhance modeling of early placenta development, trophoblast functions, and pregnancy complications compared to traditional 2D monolayer culture and with less lot-to-lot variability than natural ECM matrices.

## ACKNOWLEDGEMENTS

We acknowledge funding through the Juvenile Diabetes Research Foundation International (1-INO-2020-915-A-N), the Arizona Biomedical Research Commission New Investigator Award, the Center for Scientific Review (S10 OD023691-01), and the National Institutes of Health (NIH) Office of the Director New Innovator Award (DP2AI169476). We thank the Regenerative Medicine and Bioimaging Facility at Arizona State University (ASU) for the use of the Leica SP8 confocal microscope system, acquired by NIH SIG Award 1 S10 OD023691-01, and the KE cores facility Flow Cytometry Core for the use of the Attune NxT, especially Adam Kindelin from the Flow Cytometry Core for sharing his expertise of flow cytometry. We acknowledge the support of the Integrated Mass Spectrometry Shared Resource at the City of Hope Comprehensive Cancer Center supported by the National Cancer Institute of the National Institutes of Health under award number P30CA33572. We thank Dr. Mana Parast from the University of California, San Diego, for kindly gifting the TSC CT 1049 cell line used in this study.

## AUTHOR CONTRIBUTIONS

Conceptualization, E.M.S. and J.D.W.; Investigation, E.M.S. and R.S.; Formal Analysis, E.M.S., N.H., and J.D.W.; Writing – Original Draft, E.M.S. and J.D.W; Writing – Review & Editing, E.M.S., N.H., R.S., P.P., and J.D.W.; Funding Acquisition, J.D.W.

## SUPPLEMENTAL INFORMATION

Table S1. Excel file containing normalized abundances too large to fit in a PDF, related to Figures 4-7.

Document S1. Figures S1-S7, Supporting Information.

