## Supplemental Figures for "Engineered 3D Hydrogel Matrices to Modulate Trophoblast Stem Cell-Derived Placental Organoid Phenotype"

**Table S1. Table of Proteomics Normalized Abundances** – available in Supporting Information

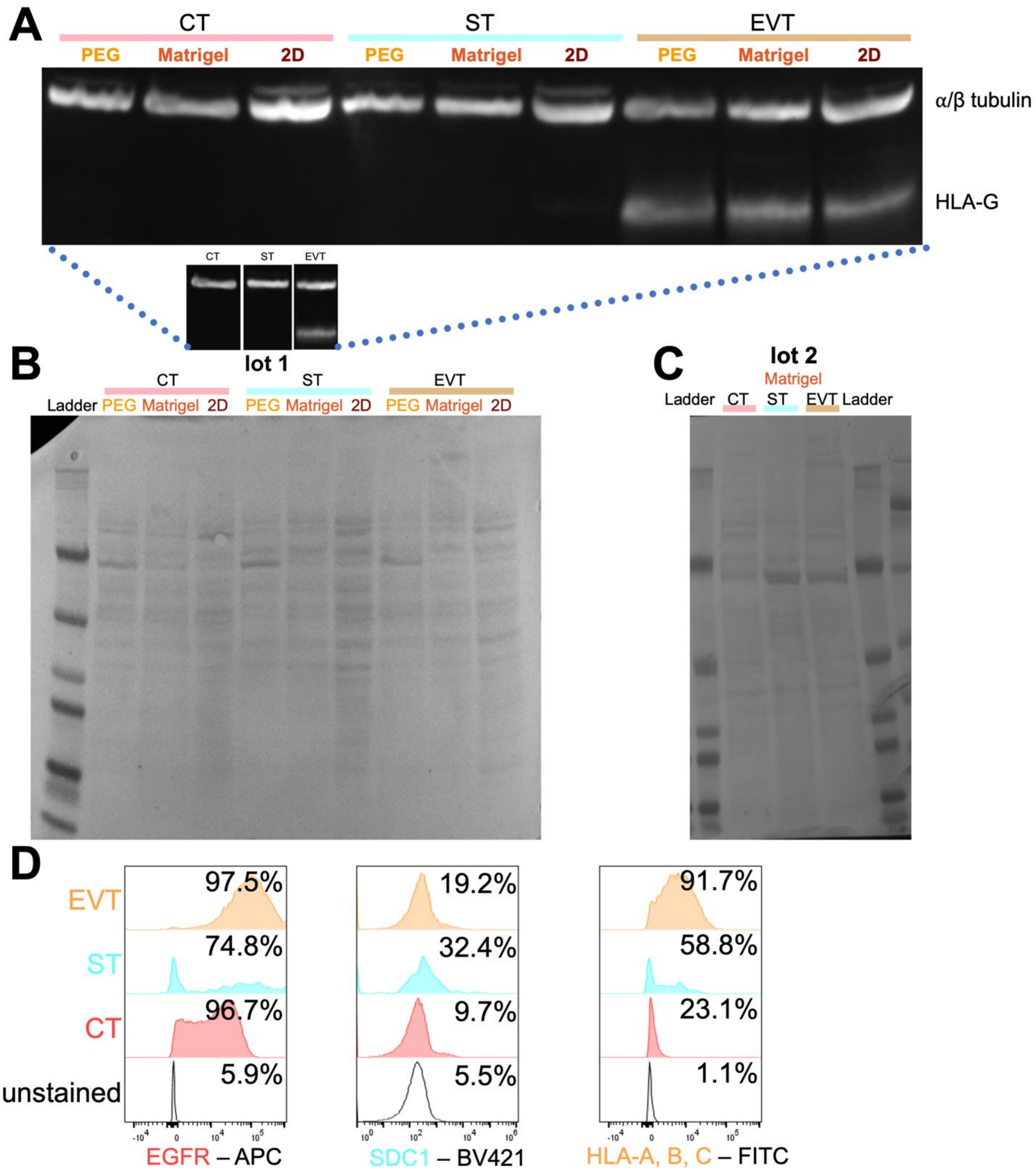


**Figure S1. Characterization of 2D TSC differentiated into CT, ST, and EVT.** (A) The full western blot from the spliced image shown in Figure 1B with corresponding (B-C) Ponceau S total protein staining of (B) lot 1 and (C) lot 2. (D) Histograms from flow cytometry of CT, ST, and EVT differentiated cells on day 6 of EGFR, SDC1, and HLA-A, B, C after gating out debris and gating on single, live events.

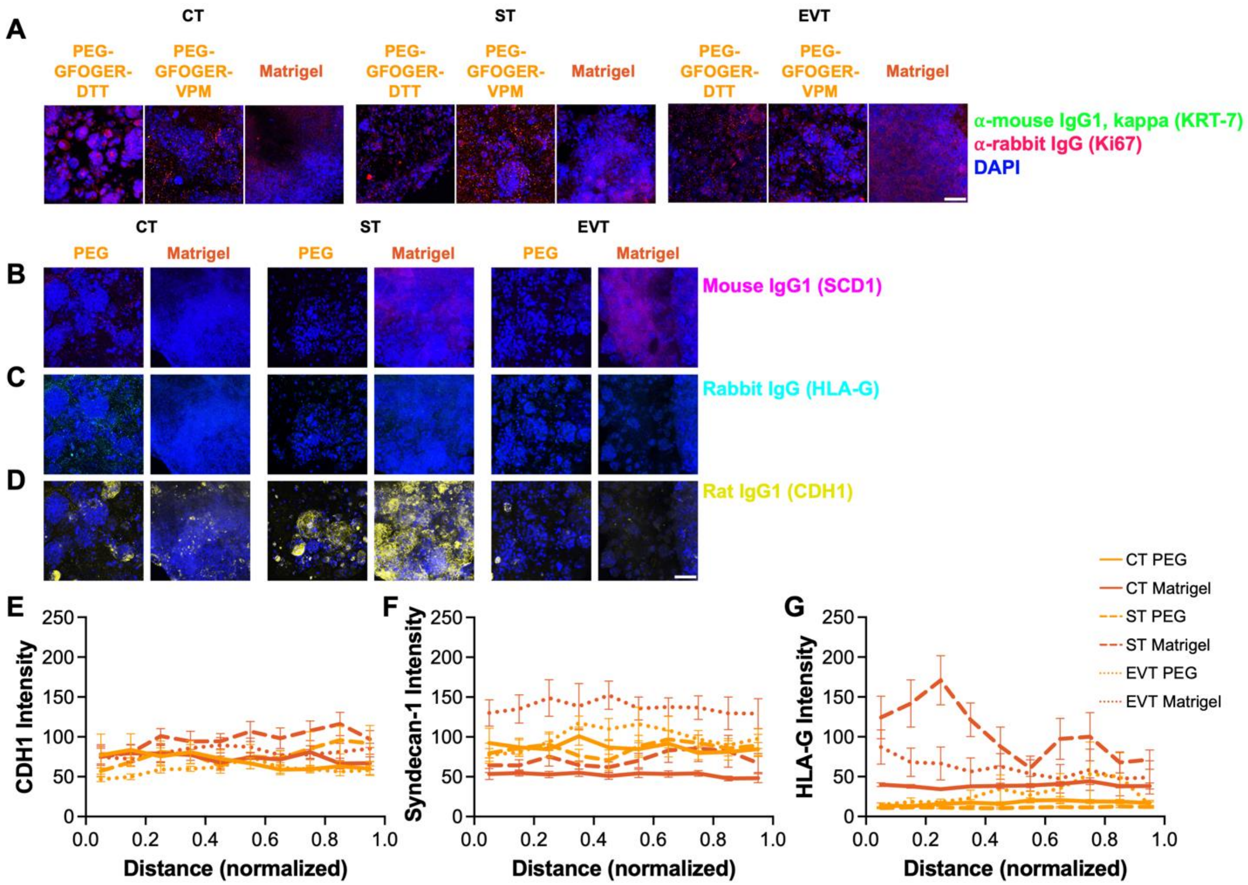


**Figure S2. Cytohistochemistry and analysis of CT, ST, and EVT differentiated from TSC.** (A-D) Isotype controls of (A) KRT7 and Ki67 staining from Figure 1C and (B) SDC1, (C) HLA-G, and (D) CDH1 from Figure 5L and (E-G) image analysis of organoids from Figure 5L. Scale bar = 200 µm. Data are shown as mean ± SD.

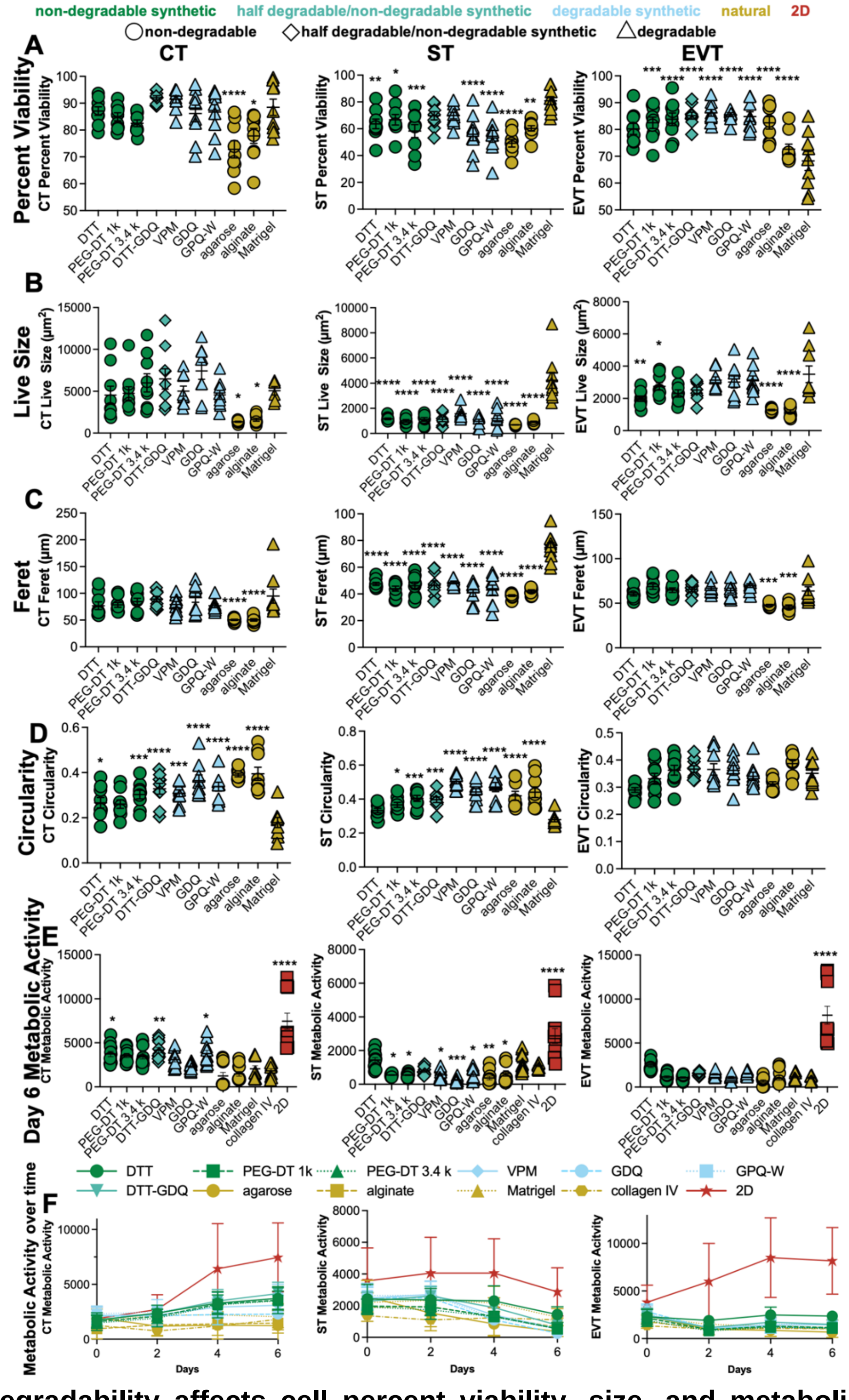


**Figure S3. Degradability affects cell percent viability, size, and metabolic activity.** (A) Percent viability, (B) live size, (C) diameter (Feret), and (D) circularity of analysis from viability images (representative images shown in Figure 2B). (E) Raw values from alamarBlue metabolic activity on day 6 and (F) over time for CT, ST, and EVT. n=12 from 3 independent experiments. Data are shown as mean ± (A-E) SEM or (F) SD and analyzed by ordinary one-way ANOVA with Dunnett's multiple comparisons test compared to Matrigel; * $p < 0.05$, ** $p < 0.01$, *** $p < 0.001$, **** $p < 0.0001$. Scale bar = 200 μm.

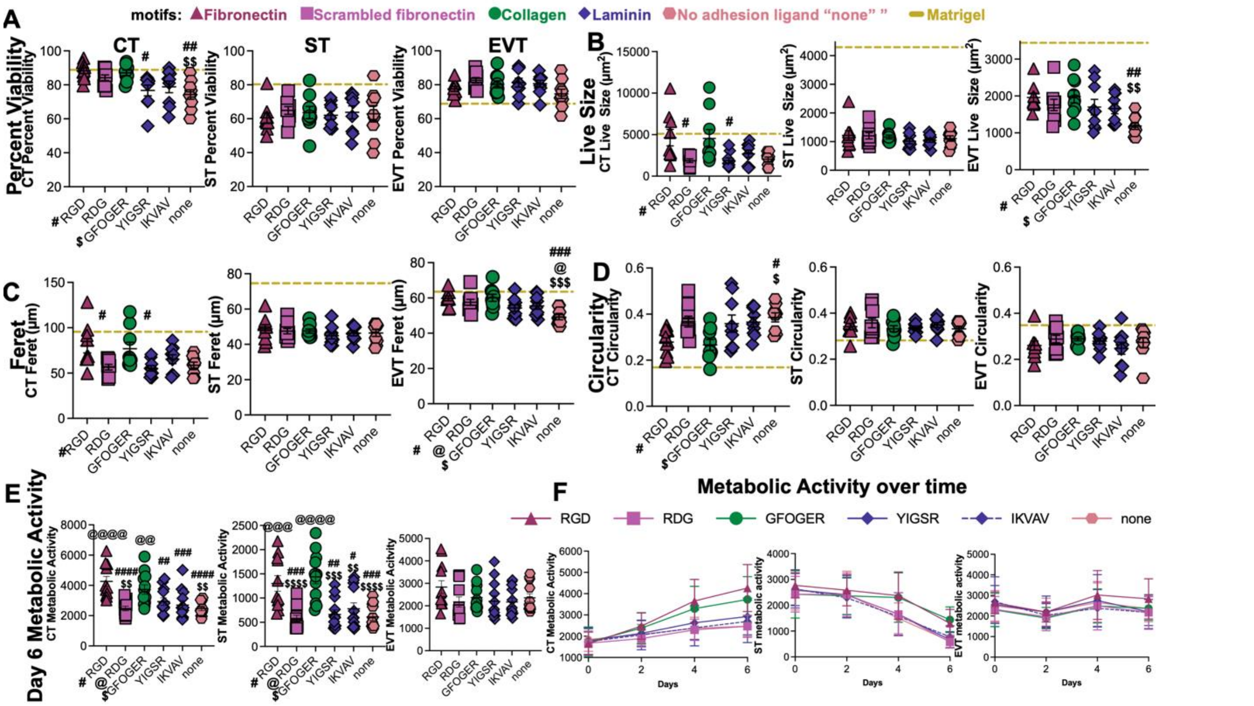


**Figure S4. Adhesion ligand presentation affects cell percent viability, size, and metabolic activity.** (A) Percent viability, (B) live size, (C) diameter (Feret), and (D) circularity of organoid analysis (representative images shown in Figure 3B). Raw values from alamarBlue metabolic activity on (E) day 6 and (F) over time for CT, ST, and EVT. n=12 from 3 independent experiments. Data are shown as mean ± (A-E) SEM or (F) SD and analyzed by ordinary one-way ANOVA with Tukey's multiple comparisons test. # compared to RGD, @ compared to RDG, and $ compared to GFOGER: * $p < 0.05$, ** $p < 0.01$, *** $p < 0.001$, **** $p < 0.0001$. Scale bar = 200 μm.

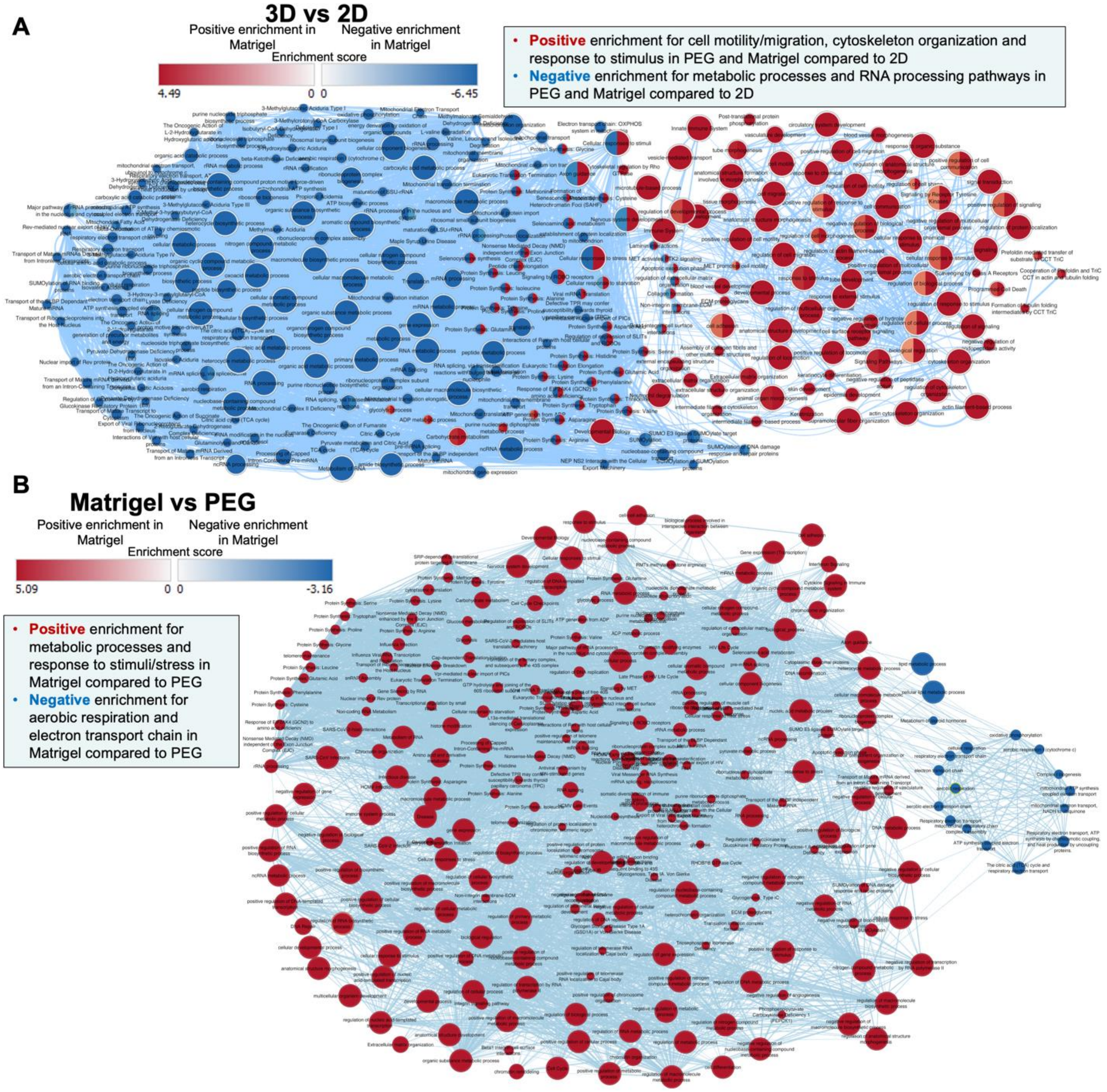


**Figure S5. Proteomics enrichment analysis reveals that culture conditions are dependent on cell processes.** (A-B) Enrichment analysis of CT, ST, and EVT differentiated in PEG-GFOGER-VPM, Matrigel, or 2D with comparisons between (A) 3D (PEG and Matrigel) versus 2D culture or (B) Matrigel versus PEG culture.

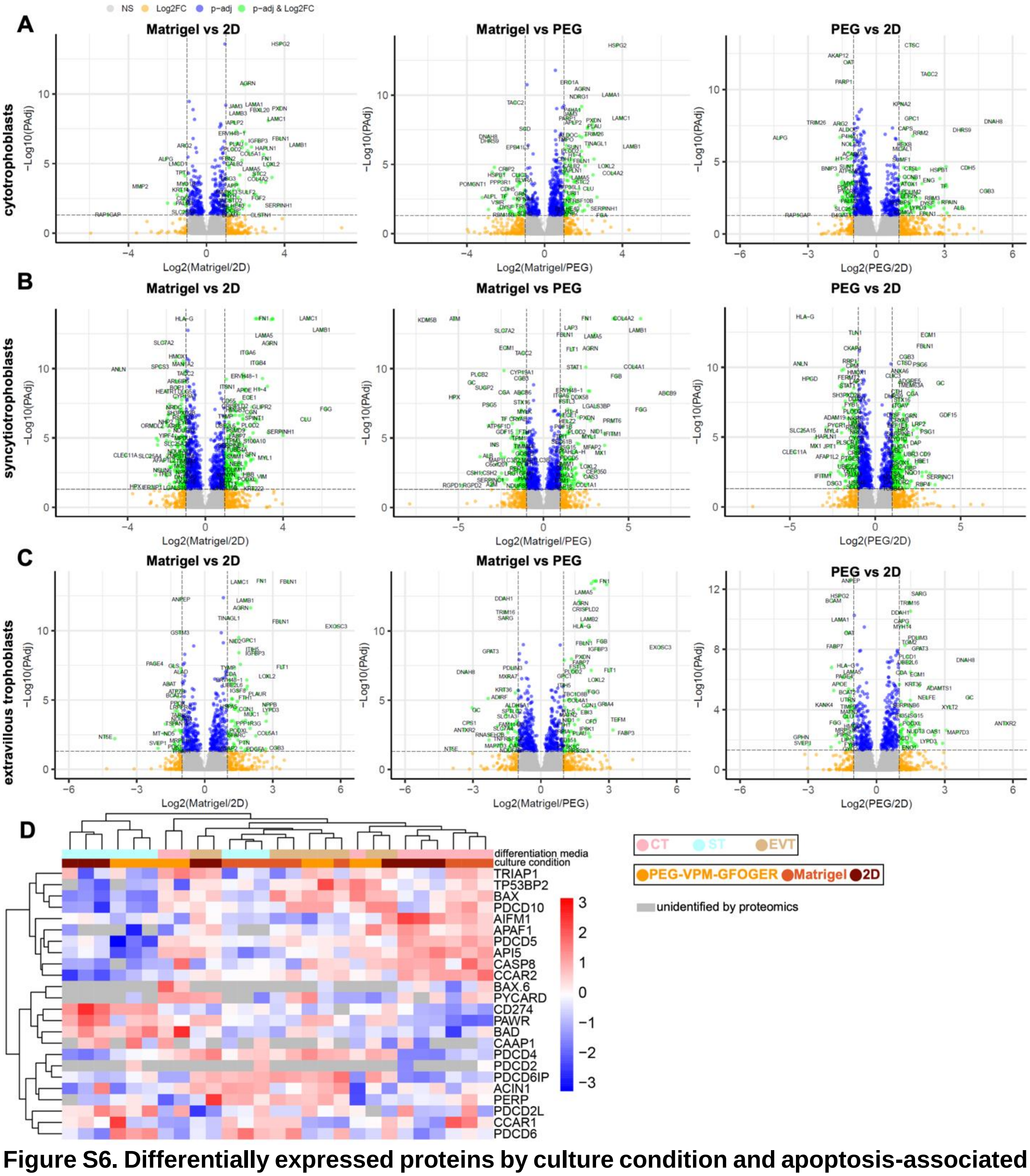


**Figure S6. Differentially expressed proteins by culture condition and apoptosis-associated proteins.** (A-C) Volcano plots of differently abundant proteins in (A) CT, (B) ST, and (C) EVT in hydrogels: Matrigel versus 2D (left), Matrigel versus PEG (middle), and PEG versus 2D (right). (D) Normalized protein abundances of apoptosis-associated proteins.

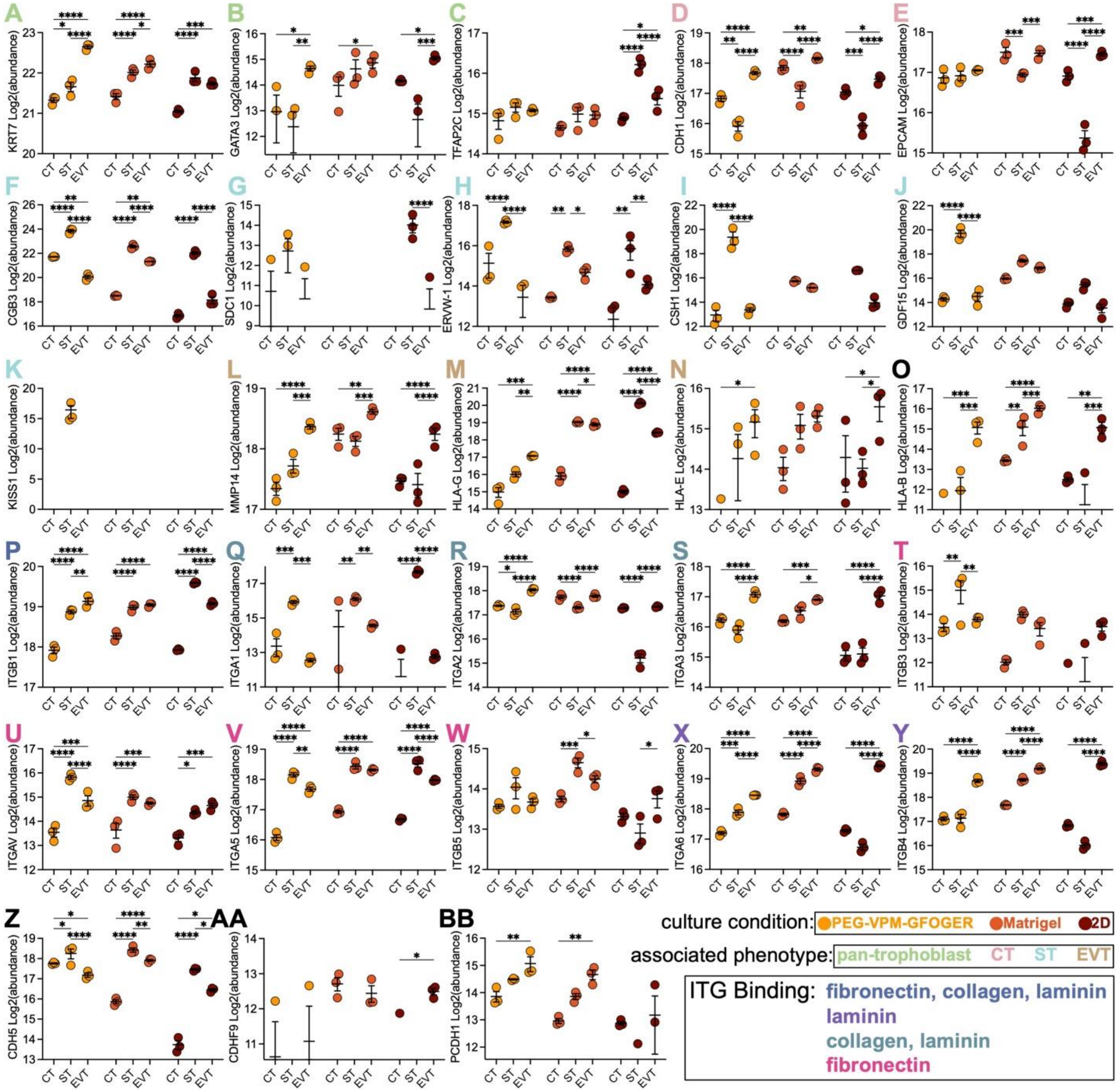


**Figure S7. Trophoblast phenotype comparison from proteomics abundances.** Protein abundances of CT, ST, and EVT (ET) cultured in PEG-GFOGER-VPM, Matrigel, or 2D with statistics shown between CT, ST, and ET grown in the same culture condition: (A) KRT7, (B) GATA3, (C) TFAP2C, (D) CDH1, (E) EPCAM, (F) CGB3, (G) SDC1, (H) ERW-1, (I) CSH1, (J) GDF15, (K) KISS1, (L) MMP12, (M) HLA-G, (N) HLA-E, (O) HLA-B, (P) ITGB1, (Q) ITGA1, (R) ITGA2, (S) ITGA3, (T) ITGB3, (U) ITGAV, (V) ITGA5, (W) ITGB5, (X) ITGA6, (Y) ITGB4, (Z) CDH5, (AA) CDFH9, and (BB) PCDH1. Data are shown as mean ± SEM and analyzed by 2-way ANOVA with Tukey's multiple comparisons test; * $p < 0.05$, ** $p < 0.01$, *** $p < 0.001$, **** $p < 0.0001$.
